# Resuscitating Sleeping Beauties: reviving a six-hundred-year-old amoeba and endosymbiont

**DOI:** 10.1101/2023.09.22.558946

**Authors:** Hasni Issam, Mamadou Lamine Tall, Matthieu Le Bailly, Philippe Colson, Anthony Levasseur, Didier Raoult, Bernard La Scola, Michel Drancourt

## Abstract

Coprolites are sources of ancient human gut microbiota and parasites. Paleomicrobiological investigation of a 14^th^-century coprolite revealed amoeba cysts that when cultured were identified as *Acanthamoeba castellanii* strain Namur. Genomic DNA sequencing indicated a 69-Mb, 57.9% GC content genome encoding 40,545 proteins, including 266 amoeba sequences sharing the best hit with amoeba-resisting microorganisms, including 170 with giant viruses. An endosymbiont we named *Coprolita marseillensis* was revealed by electron microscopy that harbored a 1-Mb genome of Rickettsiales ancestry encoding 1,738 predicted genes. Most of the amoeba genomic complement was related to biological functions giving the amoeba the capacity to survive in various ecological niches, including genes associated with anaerobic respiration and dormancy that were detected in bacteria resuscitated from permafrost, but not in controls that had not been resurrected. Investigating these 600-year-old “sleeping beauties” indicated the centuries-long survival of unicellular organisms and the mixed capability of anaerobic respiration coupled with dormancy.

## INTRODUCTION

Coprolites are fossilized feces and are a source to help decipher the composition of ancient digestive tract microbiota and identify ancient gut parasites. ^1^ While coprolites can be contaminated on the surface, the microorganisms found inside coprolites are acknowledged as those that were initially present at the time of fecal emission. ^2^ Investigating one such human coprolite dating back to the 14^th^ century, we observed cysts that yielded an *Acanthamoeba* vegetative amoeba in culture, establishing a new record of resuscitating eukaryotes from a clinical specimen. ^3^ The isolation of *Acanthamoeba* spp. from extreme environments such as artic permafrost, ^4^ frozen lakes, ^5^ and ocean sediments more than 2,500 m below the ocean surface suggests their ability to survive despite low oxygen levels. ^6^ Moreover, several studies have revealed the ability of *Acanthamoeba castellanii* to move, survive and divide at a slow rate under microaerophilic conditions. ^7^

To survive, microorganisms adopt a strategy alternating between growth and nongrowth phases. ^8^ Some microorganisms have the ability to transform into specialized cells, such as cysts, ^9^ while others, such as *Mycobacterium tuberculosis,* achieve dormancy by adapting their metabolism. ^10, 11^

The unprecedented observations reported here gave a unique opportunity to question the mechanisms underlying the long-lasting survival of a unicellular eukaryote.

## RESULTS

### Detecting amoebas in a medieval coprolite

The 14th century coprolite specimen was collected in 1996 from the interior of a closed barrel stacked at a depth of 3.80 m, used as a latrine from a Middle Age site in Namur, Belgium. The 121.4 g coprolite specimen was well preserved under anaerobic conditions and described as a mix of soil and organic matter. The analysis of 172 transmission electron microscopy images taken from 25 thin sections yielded several microorganism-like structures, mainly bacteria-like organisms. Congo red-stained cellulosic components observed by light microscopy revealed 4.1-13.5 µm large cysts whose morphology was compatible with that of amoebal cysts. ^3^ (Figure 1). One further microorganism corresponded structurally and morphologically to a small amoebal cyst (Figure 1A-E): this 3.7 µm long, 2.4 µm wide organism exhibited a typical 260 nm-thick outer layer and an inner membrane surrounding the intracellular structure. Using *Acanthamoeba-*specific primers, ^12^ PCR-based sequencing performed on the coprolite DNA extract in the presence of negative controls further yielded a sequence with 99% similarity to *Acanthamoeba castellanii* sequences (GenBank accession JF437606.1) (Table S1).

**Figure 1.**
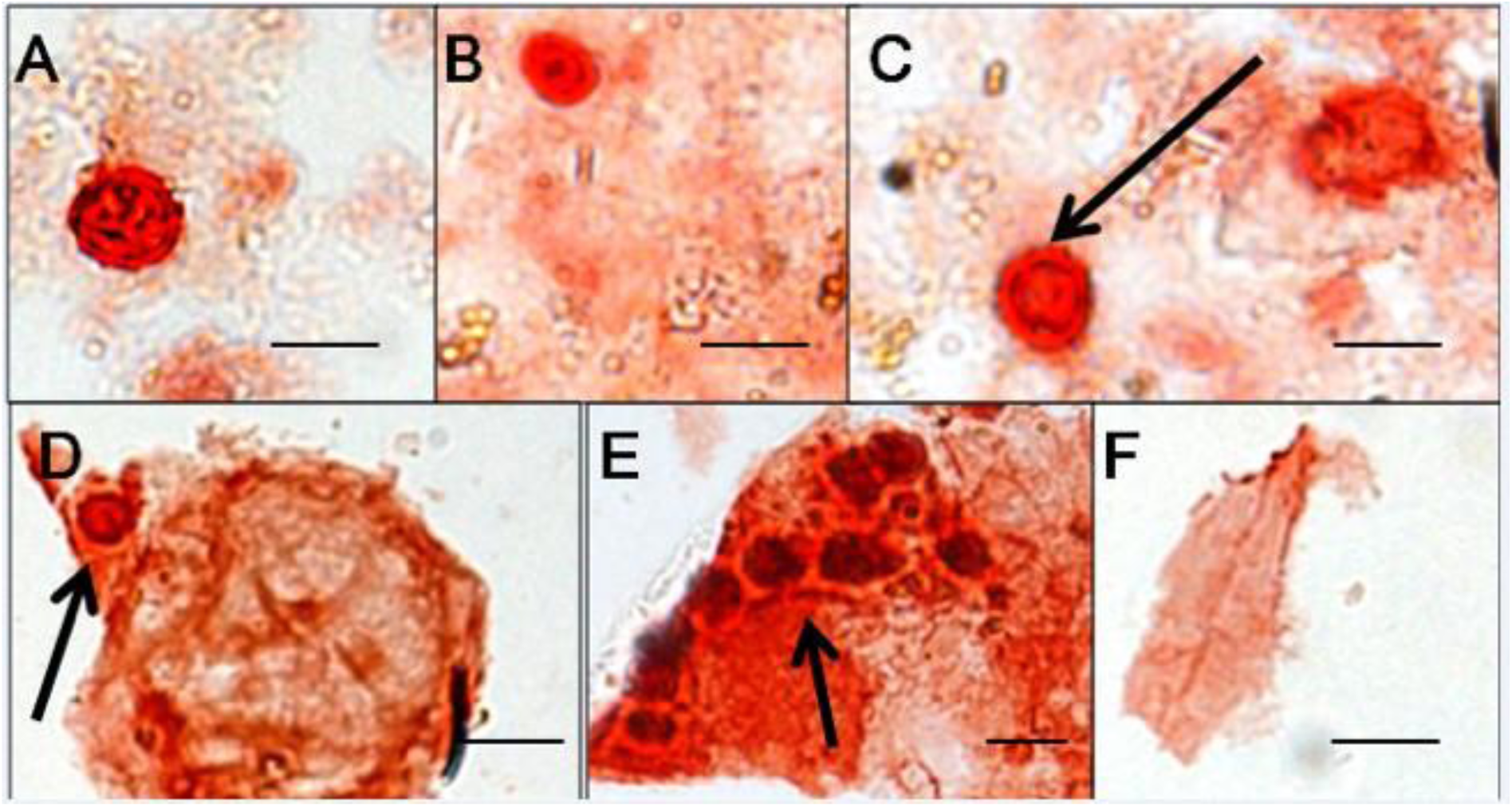
Microscopic observation of stained coprolite material. Cellulosic components were stained using Congo Red. **A-E**) Cysts, **F**) plant fiber. (Camera, Nikon digital sight DS-U1 camera; optical magnification, x 100; scale bar on the right, 10 µm)

### Culturing amoebas from a Medieval coprolite

Using laser microdissection, 25 cysts were collected and cultured under different conditions, and vegetative amoebas were observed on agar plates after seven-day incubation. All negative culture controls incorporated during experimentation remained sterile. Molecular identification by *ad hoc* PCR amplifications and sequencing indicated that this amoeba belonged to the genus *Acanthamoeba*. Cultured amoebas yielded 99% identity and 98% coverage with *A. castellanii* (GenBank accession U07413.1; KC164234.1), and this isolate, named *A. castellanii* strain Namur, was deposited in the Collection de Souches de l’Unité des Rickettsies as CSUR Q5321 (Table S1). Transmission electron microscopy further revealed an intra-amoebal bacillus with bipolar flagella that was nonmotile when observed by light microscopy (Figure 2). The 16S rRNA gene sequence (GenBank accession JQ409353.1) of the intra-amoebal bacillus yielded 100% coverage and 99% similarity with an uncultured endosymbiont (GenBank accession AF239294.1). An attempt to isolate the bacterial endosymbiont using *A. castellanii* strain Neff as a cellular host was successful (Figure S1). With respect to the source material and the location of culture, the endosymbiont was named *Coprolita marseillensis* and deposited as CSUR Q5321.

**Figure 2.**
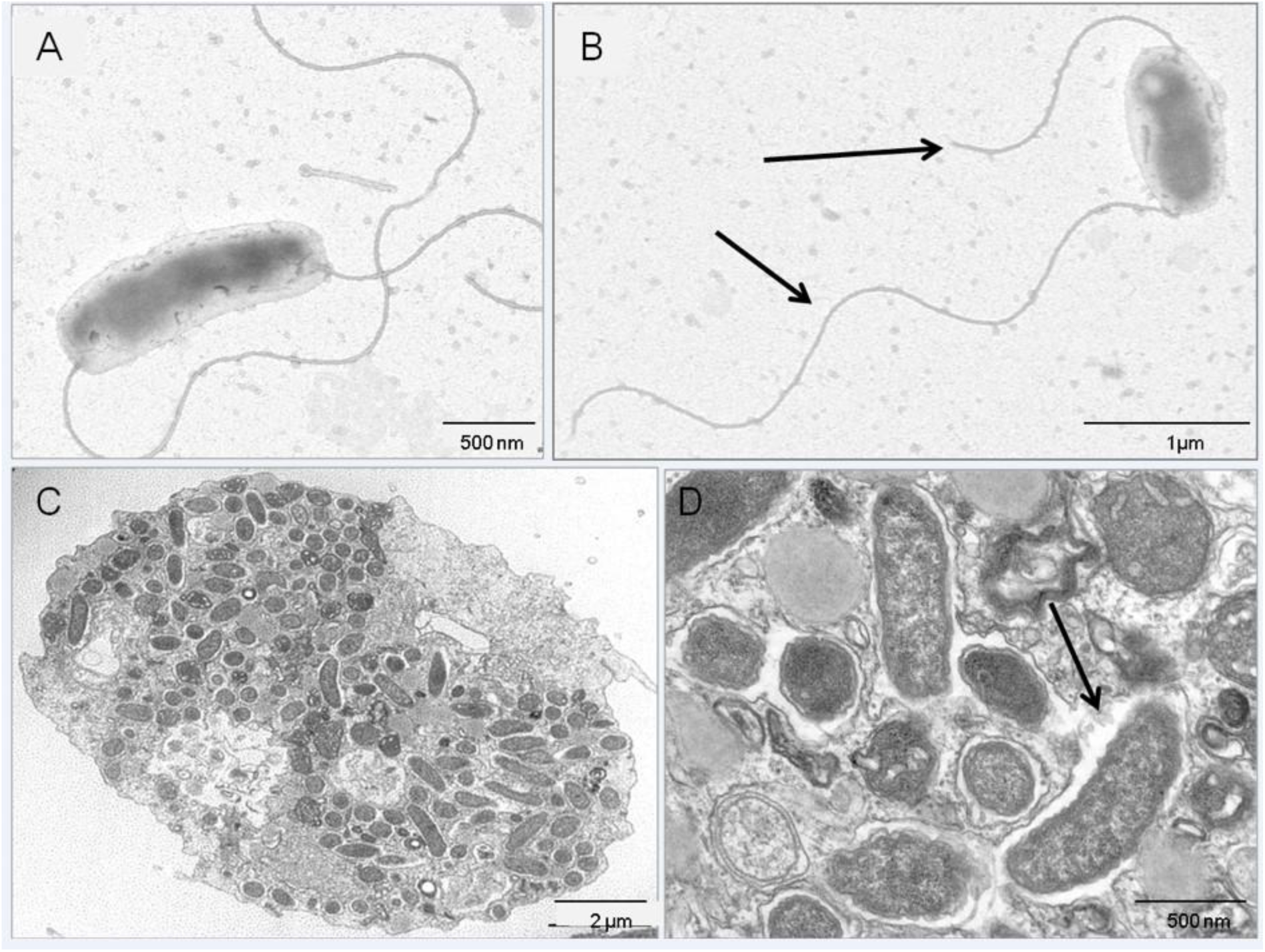
Transmission electron microscopy (TEM) observations of the *Acanthamoeba* endosymbiont. **A-B**) Observations of the endosymbiont by TEM after negative staining. **C-D**) Observations of thin layers of resin-embedded inclusion of the endosymbiont inside its host *Acanthamoeba* culture B by TEM.

### Investigating of *A. castellanii* strain Namur genome

The 69,073,894 nucleotide (nt) draft *A. castellanii* strain Namur genome encompassed 12,068 contigs, harboring a guanine-cytosine (GC) content of 57.9% (Table 1), which is similar to the GC content of *A. castellanii* (57.8%) and *A. triangularis* (58.6%). ^13, 14^ Strain Namur genome is larger than that of *A. castellanii* strain Neff (42.02 Mb) but smaller than that of *A. castellanii* ATCC 50370 (120.6 Mb). ^15, 16^ The only one 1,741-bp 18S rRNA gene copy (GenBank accession no. LR813621) exhibited 99.88% sequence similarity with *A. castellanii* strain 4CL (GenBank accession AF260724.1), phylogenetically the closest species; in agreement with 18S rRNA gene sequence-based phylogeny showing that *A. castellanii* strain Namur is most closely related to genotype T4 *Acanthamoeba,* including *A. castellanii* strain 4CL (AF260724.1) and *A. castellanii* strain ATCC 30011 (KF318462.1) (Figure S2). Of 40,545 predicted putative protein sequences, BLASTp analysis found homologous sequences for 34,603 protein sequences (85.3%), allowing the assignment of a putative function for 23,200 protein sequences (67% of proteins matched against the nr database), while 5,942 protein sequences were ORFans (14.3%) (Table S2). The taxonomic distribution of *A. castellanii* strain Namur protein-coding sequences demonstrated that 97.8% of annotated proteins were shared with eukaryotes, 1.6% with bacteria, 0.1% with archaea and 0.5% with viruses (Figure 3). Among the best BLAST hits shared with eukaryotes, we found 30,860 best hits with *A. castellanii* strain Neff, another *Acanthamoeba* with the T4 genotype. A total of 15,449 genes (38.1%) were distributed in 23 COGs. The “unknown function” group (3,150: 20.4%) was the most represented, followed by “signal transduction mechanisms” (1,950: 12.6%), “posttranslational modification, protein turnover, and chaperones” (1,592: 10.3%), “replication, recombination and repair” (991: 6.4%) and “intracellular trafficking secretion, and vesicular transport” (891: 5.7%). Among the COG functional classes, the categories “extracellular” (66: 0.4%) and “nuclear structure” (21: 0.1%) were the least represented (Figure S3). Functional assignment using the Kyoto Encyclopedia of Genes and Genomes (KEGG) database reported the presence of 7,559 sequences (19% of the genes) significantly matching *A. castellanii* protein sequences, distributed into 356 KEGG pathways. Among them, 2,904 genes were assigned to metabolic pathways and classified into 11 subcategories (Figure S3). Genes involved in carbohydrate pathways (632 genes) were the most abundant, followed by those involved in amino acid (547 genes) and lipid (461 genes) pathways.

**Figure 3.**
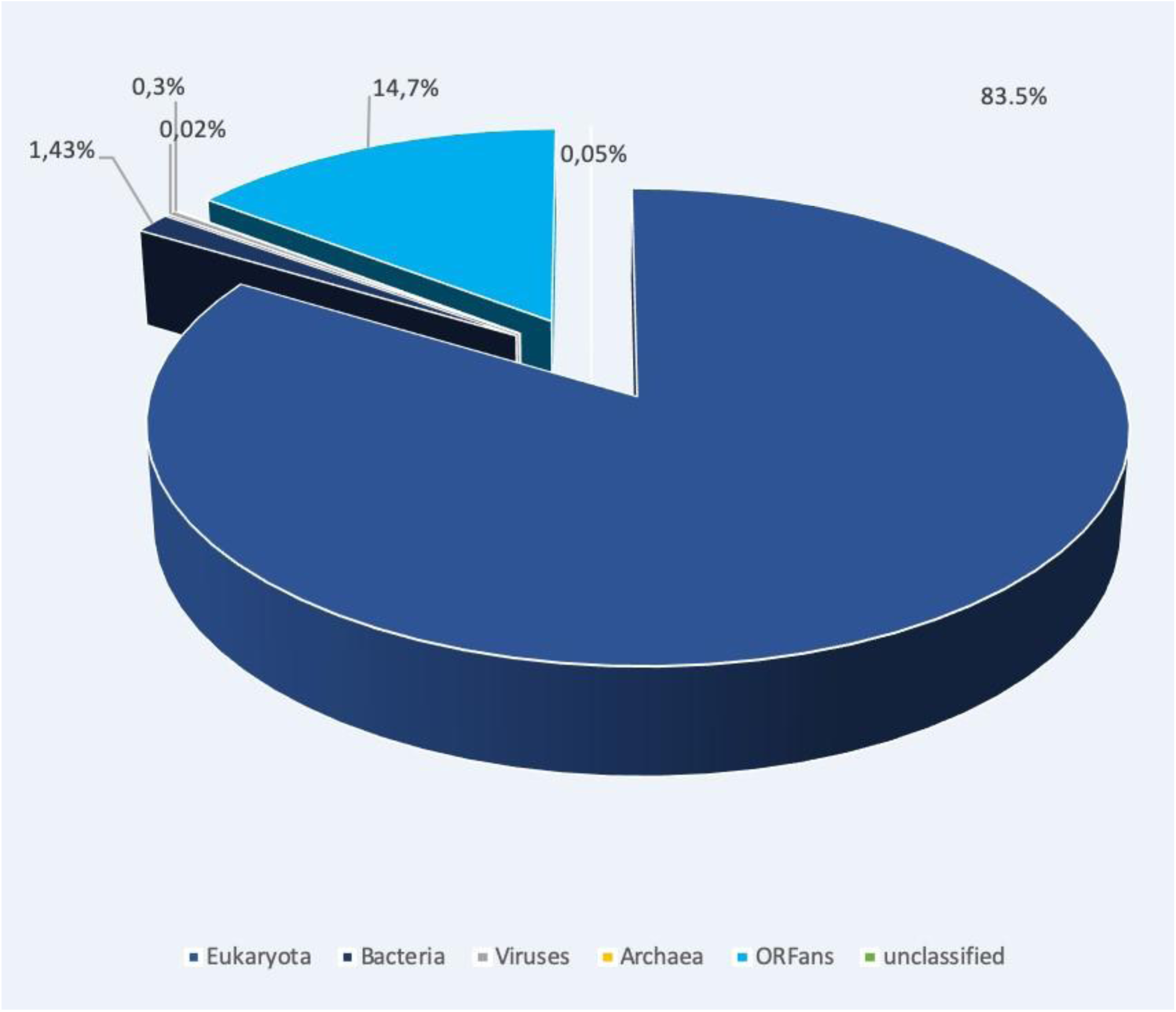
Representation of the *A. castellanii* taxonomic distribution of BLASTp best hits.

**Table 1.** Summary of the *A. castellanii* strain Namur genome. bp, base pairs; N50, 50% of the genome assembly contigs are larger than this size; N25, 25% of the genome assembly contigs are larger than this size.

| Parameter | Number |
| --- | --- |
| Haploid genome size (bp) | 69,073,894 |
| Sequence contigs (n) | 12,068 |
| GC-content (%) | 57.9 |
| Maximal scaffold size (bp) | 171,398 |
| Minimal scaffold size (bp) | 1,000 |
| Sequence | 40,545 |
| N50 | 7,685 |
| N75 | 4,393 |

### Investigation of *A. castellanii* strain Namur biology

Although *Acanthamoeba* are regarded as obligate aerobic organisms that need oxygen as the final electron acceptor to produce adenosine triphosphate (ATP), ^13^ several genes related to anaerobic respiration were identified in the *A. castellanii* strain Namur genome: we found genes encoding a pyruvate:ferredoxin oxidoreductase (PFO), an acetate:succinate CoA transferase, [FeFe]-hydrogenase (H_2_ase) and [FeFe]-H_2_ase maturase proteins HydE, HydF and HydG, homologous to sequences present in the genome of *A. castellanii* strain Neff. Indeed, homologous genes are also recovered within the genomes of free-living amoebas belonging to the family *Vahlkampfiidae,* including *Naegleria gruberi* and *Willaertia magna*. ^17, 18^ (Table S3). All these enzymes are involved in the anaerobic metabolism of protists under low oxygen conditions. ^13, 19, 20^ These genes encode factors involved in the anaerobic decarboxylation of pyruvate by PFO, producing acetyl-CoA, CO_2_ and two electrons with the concomitant reduction of NAD^+^ to NADH. Then, [FeFe]-H_2_ase transfers the electrons to 2H+ to produce hydrogen gas. The identification of an acetate:succinate CoA transferase suggested that acetyl-CoA was converted to acetate with the production of ATP (Table S3). In addition to acetate synthesis and hydrogen fermentation, we report genes involved in nitrite (nitrate reductase) and fumarate reduction (Table S3). These findings were congruent with similar observations made in some protists, especially for *A. castellanii* strain Neff, *Naegleria gruberi* and *Trichomonas vaginalis*. ^13, 17, 19, 21^ The BLASTp analysis showed that sequences related to anaerobic metabolism of *A. castellanii* were homologous to sequences belonging to strict or facultative anerobic organisms such as the protist *Mastigamoeba balamuthi*, the bacterium *Thermotoga profunda* or the archaeon *Thermoplasmatales archaeon*. These observations suggested the ability of *A. castellanii* strain Namur to switch from aerobic to anaerobic metabolism to prolong its life under anaerobic conditions such as those in coprolites. Through experimental progressive hypoxia, we analyzed the capacity of the *A. castellanii* strain Namur to switch its metabolism and produce hydrogen (Table S3). At day 0, no H_2_ or CO_2_ was detectably produced by the amoebas, whereas a large increase in the concentration of H_2_ and CO_2_ was detected after 14 days, while trophozoites were still alive in the medium and the air (O_2_ and N) quantity decreased, indicating progressive depletion of oxygen. These experimental observations correlated with *in silico* data suggesting that the amoeba had the ability to switch its metabolism to survive in an oxygen-deprived environment (Figure S4 and Table S3).

To further explore this observation, we searched for the presence of these genes within *Tomitella biformata* strain AHU 1821, a bacterium previously revived from 25,000-year-old Alaskan permafrost specimens. ^22^ We identified three sequences (pyruvate ferredoxin oxidoreductase, succinyl CoA-acetoacetate CoA transferase and FeFe-hydrogenase maturase HydE) involved in *A. castellanii* anaerobiosis that shared homology with genes of *T. biformata* strain AHU 1821 (Table S3). Out of these three genes, two shared homology with protein sequences (pyruvate ferredoxin oxidoreductase and succinyl CoA-acetoacetate CoA transferase) belonging to the genome of the plague agent *Y. pestis*, a pathogenic bacterium never revived from an ancient sample. ^23^ Therefore, we assumed that other mechanisms were active in the long-term survival of the *A. castellanii* Namur strain, analogous to mechanisms involved in dormancy of the deadly tuberculosis agent *M. tuberculosis* (Table S3). ^24^ In *M. tuberculosis*, dormancy is regulated by the 48-gene DosR operon and is activated by a progressive decrease in oxygen concentration. ^24^ We observed that 13 of these 48 genes shared homology with genes in the *A. castellanii* Namur strain (Table S3). Apart from one gene not associated with a function, our analysis reported homology with a gene encoding the hypoxic response protein, a gene of the dormancy regulon that has been reported to be highly transcribed and enhances bacterial survival under hypoxia. ^25^ This latter study reported homology with a gene encoding the DNA-binding transcriptional activator DevR/Dos, which is the principal component that regulates the DosR system. ^11^ In addition, the analysis showed homology with an alpha-crystallin protein that plays a key role in the maintenance of long-term viability during dormancy. ^26^ Among the 13 genes related to the dormancy mechanism, 5 genes shared homology with glycosyl hydrolase R, phosphoribosyl transferase Rv0571c, a polyglutamine synthesis accessory protein, universal stress protein Rv2005c and diacyglycerol O-acyltransferase sequences and the protein sequences of *T. biformata* strain AHU 1821 but not with the negative control *Y. pestis* protein sequences (Table S3). To enforce the specificity of this observation, 30 genes randomly chosen within the *M. tuberculosis* genome were used as negative controls, ^27^ and only 11 yielded homologies with *A. castellanii* protein sequences (Table S3).

Further, we found genes encoding penicillin G amidase, which is involved in the biosynthesis of penicillin derivatives such as penicillin G (Table S4). The analysis identified several genes involved in streptomycin biosynthesis (Table S4). Moreover, analysis uncovered 3 genes containing non-ribosomal peptide sequence (NRPS)/polyketide synthase (PKS) domains. These sequences seem to be involved in the metabolic pathway of epothilone and kirromycin, which are molecules of potential interest for the development of antimicrobials (Table S4). ^28, 29^ Among them, one gene (gene number 5370) encodes a potential PKS shared a best hit with a gene in the *A. castellanii* strain Neff sequence, which was annotated as an AMP-dependent synthetase and ligase (XP_004368174.1). However, the analysis of this *A. castellanii* strain Neff protein sequence domain revealed the presence of beta-ketoacyl synthase, ketoreductase, acyl transference and polyketide synthase dehydratase domains (Figure S5). Furthermore, phylogenetic reconstruction showed that the PKS sequences of *A. castellanii* strain Namur and *A. castellanii* strain Neff clustered with their homologous sequences in two fungi, *Byssochlamys spectabilis* and *Basidiobolus meristosporus* (Figure S5). According to this phylogeny, we hypothesized that the PKS sequences were transferred between amoebas and fungi. In addition to these genes related to secondary metabolites, we found multiple copies of genes encoding beta-lactamases responsible for resistance to beta-lactam antibiotics (Table S4). Numerous genes involved in the biosynthesis of natural products were identified, such as genes involved in the biosynthesis of terpene and squalene precursors. These natural products have a large range of biological properties. ^30^ The presence of several genes involved in chloroalkane and chloroalkene degradation suggested that the *A. castellanii* Namur strain had the potential to degrade molecules regarded as organic pollutants. ^31^ Furthermore, we identified several genes encoding ATP binding cassette (ABC) transporters implicated in the import and export of a large variety of substrates and potentially involved in xenobiotic detoxification.

### Investigation of horizontal gene transfers

The sequences of *A. castellanii* strain Namur were investigated using BLASTp against the nonredundant protein sequence database (National Center for Biotechnology Information). In-depth analysis of the taxonomic distribution showed that *A. castellanii* strain Namur had 266 best hit sequences with amoeba-resisting microorganisms (ARM), including giant viruses and amoeba-resisting bacteria (ARB) (Table S5). Out of 172 best hit sequences of viruses, we identified 170 *A. castellanii* protein sequences shared with giant viruses, which have the ability to infect certain eukaryotic organisms or detected during metagenome analyzes. ^32–34^ We detected homology with giant viruses belonging to *Phycodnaviridae* (104 genes: 61.2%), *Medusaviridae* (36:21.2%), *Mimiviridae* (6:3.5%), *Marseilleviridae* (6:3.5%), *Pithoviridae* (10:17%) and *Asfarviridae* (1: 0.6%) (Figure 4). The analysis indicated that the most predominant giant viruses having sequence homologies with strain Namur were pandoraviruses (68 genes: 40%), Medusavirus (36:21.2%) and Mollivirus (24:14.1%). Most of the best viral hits were assigned to proteins with important biological functions (90 genes: 53%), including ankyrin repeat protein, transposase protein (n = 4), DNA-directed RNA polymerase II subunit RPB1 (n = 1) and endonuclease V. In addition to viral homologs, we detected the presence of best hits shared in ARBs. All these bacteria have the capacity to infect and survive within free-living amoebas. ^32, 35^ The analysis reported the presence of 96 best hits in bacteria belonging to the *Chlamydiae* phylum (89 genes; 92.3%), *Pseudomonas* spp. (3 genes; 3.1%), *Acinetobacter* spp. (2 genes; 2.1%), *Mycobacterium* spp. (1 gene; 1%) and *Legionella* spp. (1 gene; 1%) (Table S5). To obtain more information on the functions of ARM protein sequences, we searched with BLAST against the COG database. The protein sequences involved in “replication, recombination and repair” (n = 17: 6.4%) were the most represented among COG categories, followed by “function unknown” (n = 13: 4.9%) and “transcription” (n = 5: 1.9%) (Table S5). The investigation of putative horizontal transfers was assessed by a phylogenetic tree based on the *A. castellanii* homologs with ARMs. Out of 266 potential sequences shared with ARMs, we were able to achieve 193 phylogenetic reconstructions (Figure 5 and Table S5). The protein sequences with insufficient numbers of hits did not allow us to examine the potential horizontal transfers, as was the case for 73 sequences. Horizontal gene transfers were confirmed by phylogenetic investigation for 155 (58.3%) protein sequences, including 108 and 47 protein sequences that had best hits with giant viruses and ARBs, respectively. The analysis based on BLASTp comparison showed that the DNA polymerase sequence of *A. castellanii* (gene 15,991) shared homologs with giant viruses, including Mollivirus and 7 Pandoravirus strains. Furthermore, phylogenetic reconstruction reported clustering of these DNA polymerase homologs. This result suggested potential horizontal gene transfer from *A. castellanii* to giant viruses (Figure 6).

**Figure 4:**
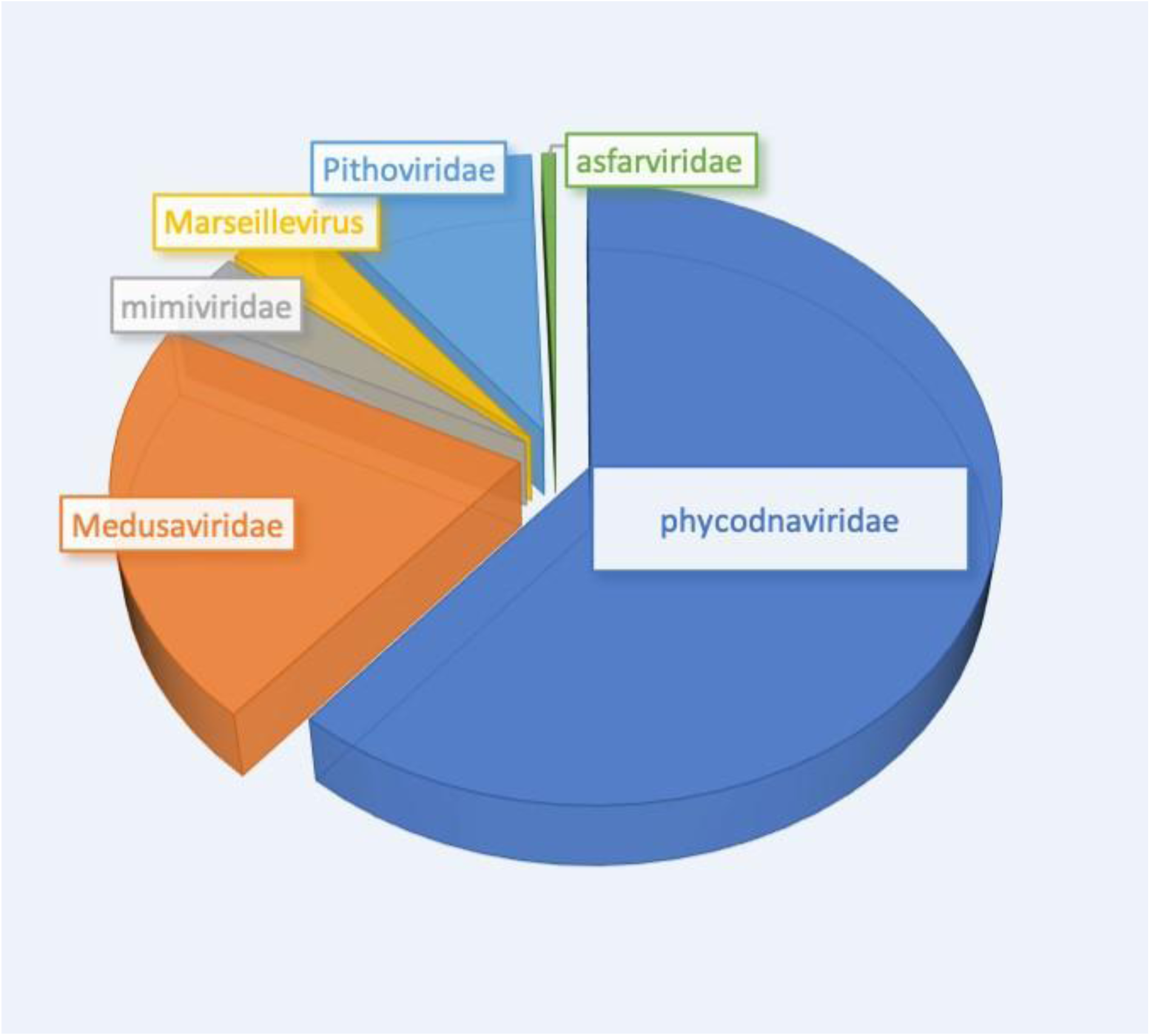
Representation of the giant virus family best matching the *A. castellanii* protein sequences.

**Figure 5:**
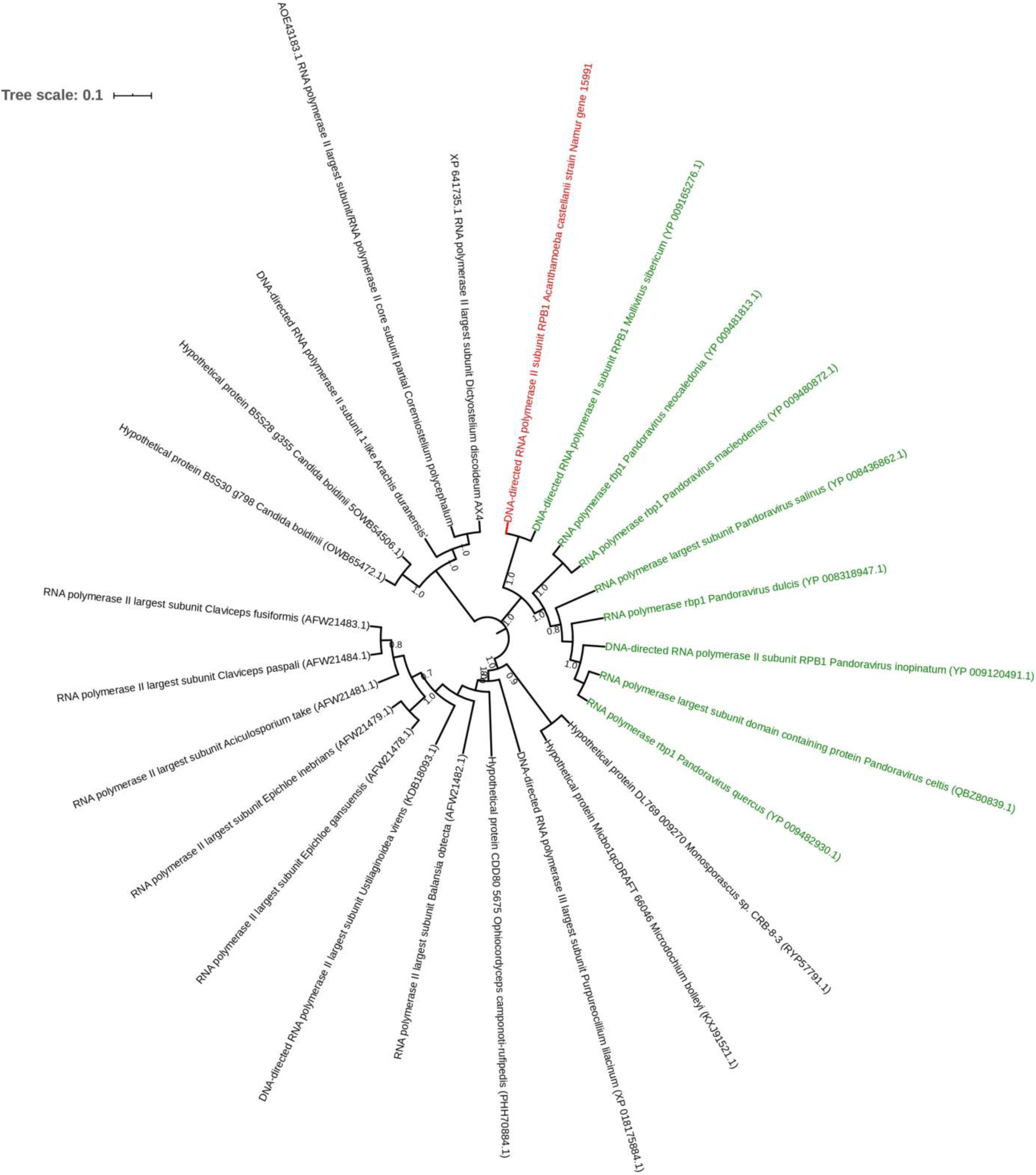
Investigation of lateral gene transfers. Phylogenetic trees for four *A. castellanii* protein sequences of putative ARM origin. The trees were constructed using the maximum-likelihood method based on the hydrolase, metallophosphatase, TPR repeat protein and hypothetical protein sequences of *A. castellanii* strain Namur. **A**) In red, hydrolase of *A. castellanii* strain Namur; in blue, the closest homologs from ARM (Pacmanvirus) and in black, homologs from other organisms. **B**) In red, metallophosphatase of *A. castellanii*; in blue, homologs from ARM (Pandoravirus strains); in black, homologs from other organisms. **C**) In red, TPR repeat protein of *A. castellanii* strain Namur; in blue, homologs from ARM (*Acinetobacter calcoaceticus*); in black, homologs from other organisms. **D**) In red, hypothetical protein of *A. castellanii* strain Namur; in green, homologs from ARM (*Candidatus* Proteochlamidya amoebophilus); in blue, homologs from *Acanthamoeba castellanii* strain Neff; in black, homologs from other organisms.

**Figure 6:**
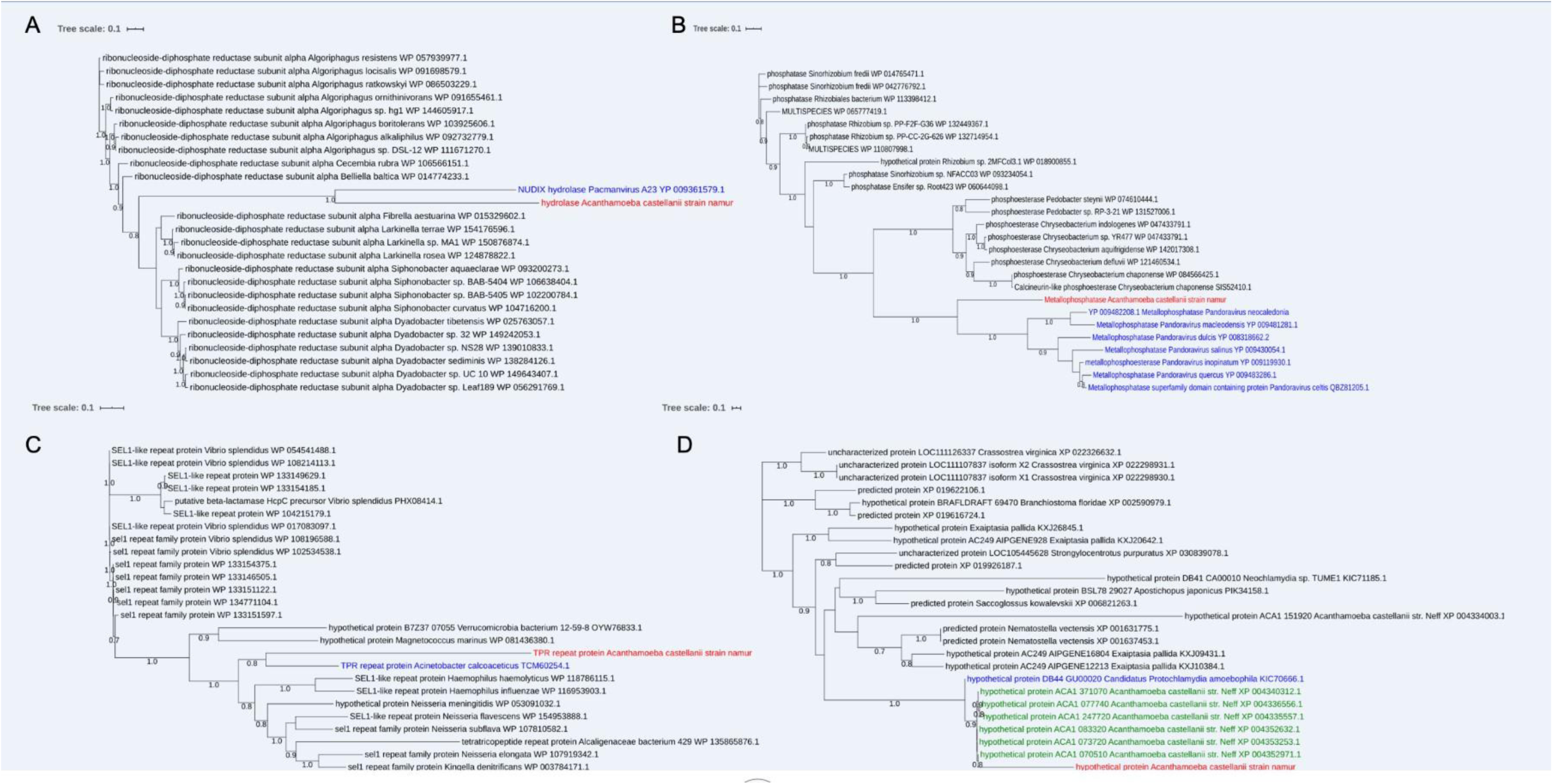
Investigation of lateral gene transfers. Phylogenetic trees for DNA-directed RNA polymerase II of the *A. castellanii* protein. The trees were constructed using the maximum-likelihood method based on the DNA-directed RNA polymerase II protein of *A. castellanii* strain Namur. In red, DNA-directed RNA polymerase II of *A. castellanii* strain Namur; in green, the closest homologs from ARM (giant viruses); in black, homologs from other organisms.

### Endosymbiont genome

The 1,928,597-bp genome (81 scaffolds) of *C. marseillensis* exhibited a 34.3% GC content. Of the 1,738 predicted genes, 1,699 were protein-coding genes, and 39 were RNAs (3 rRNAs and 36 tRNAs). A total of 1,044 genes were assigned a putative function, and 658 genes were annotated as hypothetical proteins (Table S6). The COG investigation assigned COG classifications to 1,243 genes (71.2%) and distributed them in 20 COG groups (Table S6). The cluster for “Function unknown” (206: 16.5%) was the largest, followed by “translation, ribosomal structure and biogenesis” (152: 12.2%), “replication, recombination and repair” (148: 11.9%) and “energy production and conversion” (77: 6.2%). Among the COG functional classes, the categories “cytoskeleton” (4: 0.3%) and “RNA processing and modification” (2: 0.1%) were less represented (Figure 2). Comparing COG annotation in *C. marseillensis* and phylogenetically closely related organisms revealed similar COG distributions except for the case of *Rickettsia typhi* (Figure S6). A 16S rRNA gene sequence-based phylogenetic tree clustered *C. marseillensis* and *Lyticum flagellatum,* while phylogenetic reconstruction based on draft genomic data indicated that *C. marseillensis* was more closely related to *Rickettsia raoultii* (Figure S7). Comparing the genomic content of *C. marseillensis* with that of phylogenetically closely related organisms yielded OrthoANI values ranging from 64.66% with *Rickettsia sibirica* to 99.17% with *Rickettsia parkeri* (Figure S7). Furthermore, we found 10 *C. marseillensis* genes sharing homology with protein sequences involved in *M. tuberculosis* dormancy (Table S3). Among these genes, four shared homologies either with *A. castellanii* (alpha-crystallin and uncharacterized protein Rv2030c) or with *T. biformata* (ferredoxin and uncharacterized protein Rv1734c) genes, while there were no homologies with *Y. pestis* genes (Table S3). The phosphoribosyl transferase Rv0571c gene is the sole gene shared between *A. castellanii*, *T. biformata* and *Candidatus* C. marseillensis and is absent in the negative control *Y. pestis* (Table S3). Among 30 genes randomly chosen as negative controls within the *M. tuberculosis* genome, only 5 shared homologies with *C. marseillensis* gene sequences (Table S3).

## DISCUSSION

Reviving a 600-year-old amoeba and its Rickettsiales endosymbiont provided a unique opportunity to question the mechanisms underlying century-long survival of unicellular life after previous resuscitation from permafrost of free-living amoebas, ^4^ ciliates or giant viruses.^33, 34^

Oxygen depletion is known to promote amoeba encystment. ^7^ Accordingly, the *A. castellanii* Namur strain genome encodes a set of genes allowing the amoeba to produce ATP in the absence of oxygen, some of which are also present in the endosymbiont. Additionally, hypoxia is known to regulate an operon that mediates dormancy in *M. tuberculosis*, a condition preventing the death of the pathogen during a progressive decrease in oxygen. ^11^ We discovered that some *M. tuberculosis* dormancy genes had homology with *A. castellanii* Namur strain genes and the genes of its endosymbiont, suggesting that progressive anaerobia induced dormancy in addition to inducing anaerobic respiration to promote the long-term survival of both organisms. In addition, under conditions of progressive hypoxia, *A. castellanii* strain Namur and its endosymbiont were able to produce metabolites specific to the anaerobic metabolism of protists.

In the presence of oxygen, sugar oxidation generates considerable amounts of energy and the formation of reactive oxygen species (ROS), which are extremely harmful to organisms.^36^ The accumulation of oxidative damage in proteins causes protein damage and leads to the degeneration of all cellular functions. ^37^ Alternative anaerobic metabolisms in the *A. castellanii* Namur strain may have enabled the amoeba to decrease deleterious protein oxidation coupled with dormancy. Our analyses revealed the presence of genes related to dormancy in a 25,000-year-old amoeba resuscitated from permafrost. A list of specific genes related to the dormancy mechanism was only detected within organisms resuscitated from ancient samples and not from the negative control. These observations suggested that the oxygen-depletion stress response and dormancy mechanism were essential for the long-term survival of the amoeba and its endosymbiont.

The *A. castellanii* Namur strain formed an ecosystem within itself, promoting horizontal gene transfers between viruses, bacteria and fungal organisms; most of the transferred candidates encoded metabolic functions, indicating the contribution of lateral transfers in shaping amoebal biology and diversification, as previously reported. ^13^ In particular, the *A. castellanii* strain Namur genome encodes secondary metabolites involved in the biosynthesis of polyketides, streptomycin or gentamycin, and phylogenetic reconstructions yielded indications of lateral gene transfers for PKS genes between amoebas and fungi. ^14^

The present study revealed that genes encoding anaerobic respiration coupled with dormancy probably supported the long-term survival of two unicellular “sleeping beauties” during oxygen deprivation, inviting future explorations of this condition in pluricellular organisms.

## MATERIALS AND METHODS

### Contamination prevention in holding ancient coprolite

The coprolite was handled only in a positive pressure room with isolated ventilation under strict aseptic conditions. Workbenches were stringently disinfected using absolute ethanol and UV irradiation for at least 30 min. Non-disposable instruments were autoclaved. Reagents and chemicals were obtained from new stocks aliquoted into sterile, single-use tubes and immediately discarded after use. The external portion of the coprolite was aseptically removed, and only the internal portion was used in this study. DNA extraction, PCR and post-PCR experiments were performed in separate rooms in isolated work areas. Positive controls were strictly avoided. Negative controls were used in a 1:4 control: specimen ratio. ^38, 39^

### Microscopy and laser microdissection

The cellulosic material of the coprolite was stained using Congo red. ^40^ Laser dissection of the cysts was then performed using silicone membrane-coated slides (Molecular Machines & Industries, Aartselaar, Belgium), a Nikon ECLIPS TE2000-U microscope (Nikon Instruments, Champigny sur Marne, France) and an MMI CellCut laser and its controlling program MMI cell tool (Molecular Machines & Industries).

### Culture

### Culturing amoebas

A 500 mg portion of the interior region of the Namur coprolite was solubilized in 1 mL of sterile Page’s Amoeba Saline medium (PAS medium : 2 mM NaCl, 16 μM MgSO4, 27.2 μM CaCl2, 1 mM Na2HPO4, 1 mM KH2PO4), mixed with 50 µL of amphotericin B (10 µg/mL). For culturing, nonnutritive agar plates were overcoated with a thin film of living *Enterobacter aerogenes* cells. Then, 50 µL of the coprolite suspension was immediately plated (Figure S1). Negative controls consisted of PAS without coprolite material, added at a ratio of 1:1. The cultures were incubated at 28° C and observed daily by light microscopy. To axenize the cultivated *Acanthamoeba* spp., subculture was carried out using a single clone each time, first on nonnutritive agar plates with living *E. aerogenes* cells, then on three successive nonnutritive agar plates with UV-irradiated dead *E. aerogenes* cells and finally in sterile liquid proteose peptone-yeast-glucose (PYG) growth media. ^41^ When visible growth was observed, the PYG-growth medium was changed. Strain Namur was deposited in the CSUR under number CSUR Q5320.

### Culturing the endosymbiont

To culture the amoeba endosymbiont, 7 g of coprolite material solubilized in 25 mL of ddH2O were incubated overnight at 4° C. The supernatant was filtered using 0.1 µm Millipore IsoporeTM membrane filters (Millipore, Molsheim, France) and subsequently used for amoeba coculturing as previously described. ^35^ The supernatant of the culture was then used for transmission electron microscopy observations after negative staining using a 1.5% molybdate solution or after inclusion in acryl resin. The supernatant of the culture was further used for DNA extraction. The endosymbiont was deposited in the CSUR under number CSUR Q5321.

### Culture of the endosymbiont on another host

Strain Namur was cultivated in 20 mL of starvation medium at a concentration of 1.10^6^ amoebas/mL for 15 days at 28° C, lysed, and the supernatant was centrifuged at 2,000 x g for 10 min. ^42^ We inoculated 500 µL of the supernatant into a 25 cm^3^ flask containing *A. castellanii* strain Neff (ATCC 30100) in starvation medium and incubated at 28° C for 15 days. The screening of endosymbiont was performed by both examination under an inverted microscope and by scanning electronic microscopy: a suspension of supernatant was immersed in a 2.5% glutaraldehyde fixative solution, a drop was deposited onto a slide, gently washed with water, air-dried, and examined under an Emission Scanning Electron Microscope SU5000 (Hitachi, Tokyo, Japan). For the positive and negative controls, we examined the supernatant of the strain Namur culture and the strain Neff culture, respectively, both cultivated without the endosymbiont.

### Molecular identifications

Total DNA was extracted from the coprolite as previously described. ^43, 44^ Phenol-chloroform DNA protocol was used to extract total DNA from the amoebal cysts collected via microdissection. DNA was further extracted from cultured *Acanthamoeba* spp. following the supplier’s instructions for QIAamp DNA Extraction Mini Kit (QIAGEN, Courtaboeuf, France). Each extraction included several extraction blanks consisting of sterile water. *Ad hoc* PCR protocols were applied to amplify the 18S rRNA gene of *Acanthamoeba* spp. ^12^ Amplicons were generated from the DNA extract from the coprolite using suicide PCR amplification and then from cultured *Acanthamoeba* spp. ^45^ PCR was performed in 50 µL containing 1x PCR buffer, 2 µL of 25 mM MgCl_2_, 200 µM of each dNTP, 1 μL of 10 pM of each primer, 31.15 µL of ddH2O, 1 unit of HotStart Taq Polymerase (Invitrogen, Villebon-sur-Yvette, France) and 57-112 ng of DNA extract. The PCR steps included an initial incubation at 95° C for 15 min; 38 cycles of denaturation at 95° C for 1 min, annealing for 45 sec at the corresponding primer annealing temperatures, and elongation at 72° C for 90 sec; and a final elongation at 72° C for 10 min; all these steps were performed in a Gene Amp PCR System 2700 ABI Thermocycler (Applied Biosystems, Villebon-sur-Yvette, France). Resulting sequences were assembled using ChromasPro software and compared with reference sequences in the GenBank database using NCBI BLAST searches. To generate the genotype from the *Rns* profiles, the sequences were aligned and compared to the *Acanthamoeba* genotype names stored in the NCBI GenBank database. ^46^ To identify the endosymbiont, the supernatant of the culture was used for DNA extraction as described above, and the 16S rRNA gene was sequenced and compared with 16S rRNA sequences of the GenBank database using NCBI BLAST searches.

### Hydrogen concentration measurements

When the amoeba formed a trophozoite monolayer, the latter were detached from the culture flask and harvested by centrifugation at 1,000 g for 10 min followed by two washing steps using PYG. Amoeba resuspended in PYG were quantified using a KOVA® slide cell (KOVA International, CA, USA) dyed with Trypan blue (Sigma-Aldrich, Lyon, France) and amoeba concentration was adjusted to 2.10^5^ cells/mL in PYG. To put the amoeba under progressive oxygen depletion, we plated 10 mL of amoeba solution in Hungate tubes tightly sealed with rubber stoppers and incubated at 28° C for 14 day. ^47^ On days 0 and 14, we measured the levels of hydrogen, carbon dioxide and a mixture of nitrogen and oxygen using a Clarus 580 gas chromatography system (Perkin Elmer, Villebon-sur-Yvette, France) as previously described. ^48^ Calculations integrated considered gas densities at 25° C (Gas Encyclopedia website, Air Liquide), hydrogen quantities were expressed in µmol/L or parts per million (ppm) in the volume of sampled air, later being calculated using a converter available on the internet (Lenntech, The Netherlands). The amoebas were quantified (days 0 and 14) using a KOVA® slide cell counting chamber (KOVA International, CA, USA) with Trypan blue staining (Life Technologies, Carlsbad, CA, United States). These experiments were performed in triplicate.

### Genomic study

### Genome annotation and analysis of *Acanthamoeba castellanii* strain Namur

After 72 hours, *A. castellanii* strain Namur amoeba detached from the trophozoite monolayer were resuspended in PAS, and DNA extracted using the QIAamp DNA Extraction Mini Kit (QIAGEN), purified using Agencourt AMPure XP beads (1.8x ratio, Beckman Coulter Inc., Fullerton, United States) was quantified using a Qubit assay with a high sensitivity kit (Life Technologies) to 34.4 ng/µL. The mate pair library was prepared with 1.51µg of genomic DNA using the Nextera mate pair Illumina guide. The genomic DNA sample was simultaneously fragmented and tagged with a mate pair junction adapter. The fragmentation profile was validated on an Agilent 2100 Bioanalyzer (Agilent Technologies Inc., Santa Clara, CA, USA) with a DNA 7500 LabChip. The DNA fragments ranged from 1.6 kb to 13 kb with mean sizes of 2.57 and 6.99 kb, respectively. No size selection was performed, and 188 and 721 ng of tagged fragments were circularized, respectively. The circularized DNA was mechanically sheared to small fragments with optimal sizes of 1,022 and 1,380 bp, respectively, on the Covaris device S2 in T6 tubes (Covaris, Woburn, MA, USA). The library profile was visualized using a High Sensitivity Bioanalyzer LabChip (Agilent Technologies Inc.), and the final concentrations of the libraries were measured at 3.3 and 6.4 nmol/L, respectively. The libraries were then normalized to 2 nM and pooled. After a denaturation step and dilution at 15 pM, the pooled libraries were loaded into the reagent cartridge and then onto the instrument along with the flow cell. Automated cluster generation and sequencing runs were performed in a single 39-h run of 2 × 151 bp. Total information of the two flow cells at 1.2 and 9.8 Gb was obtained from a 111,000 and 913,000 cluster density per mm^2^ with cluster passing quality control filters of 96.3 and 95%. Within these runs, the index representation for *A. castellanii* strain Namur was determined at 7.13 and 84.20%. A total of 14,210,902 paired reads were obtained from the two runs. Raw data from DNA sequencing were checked using FastQC software (https://www.bioinformatics.babraham.ac.uk/projects/fastqc/). Trimmomatic software was used to trim the raw data remove adaptors and reads with an average quality above 28. ^49^ All trimmed reads were assembled *de novo* using the Spades program. ^50^ The contigs with sizes under 1,000 bp were removed. We conducted BLASTn searches against local databases with the megablast option to identify and remove scaffolds that were likely to have originated from bacterial or viral contaminants. ^51^ Then, the 18S rRNA sequence of *A. castellanii* strain Namur (CAIJLO010000000; LR813621) was identified by BLASTn comparison between the *A. castellanii* genome assembly and the 18S rRNA sequence of *A. castellanii* strain ATCC 30011 18S rRNA (KF318462.1) available in the NCBI GenBank nucleotide sequence database (nt). Multiple sequence alignment was carried out by using Muscle software. ^52^ Finally, a phylogenetic analysis of the nucleotide sequences was performed using MEGA version 7 and the maximum likelihood (ML) algorithm, with 1,000 bootstrap replicates. ^53^ Open reading frame (ORF) prediction was performed using AUGUSTUS, a software optimized for eukaryotic genomes by comparison with the NCBI nonredundant sequence database (nr) and COG database. ^46, 54, 55^ First, the function of protein sequences was identified by BLASTp against the nr database with an e-value cutoff of 1e-03. ^51^ COG annotation was performed using EggNOG with standard parameters and diamond as the mapping mode. ^56^ The metabolic pathways and biological functions of the genes were analyzed by comparison of *A. castellanii* protein sequences against the Kyoto Encyclopedia of Genes and Genomes using BlastKOALA online software with standard parameters. ^57^ Further investigation of anaerobic respiration was performed by a BLASTp comparison of *A. castellanii* protein sequences against anaerobic protein sequences and EST of *A. castellanii* Neff genome. ^21^ Each protein sequence related to anaerobic metabolism was compared by BLASTp against the protein sequences of *Y. pestis* strain Antiqua (ASM83482v1) and *T. biformata* strain AHU 1821 (BAVQ00000000.1). To obtain the protein sequence set of *T. biformata* strain AHU 1821, we predicted the proteins using GenMarkS online. ^58^ To investigate the anaerobic mechanism, *A. castellanii* protein sequences were compared by BLASTp against the protein sequences of the DosR operon belonging to *M. tuberculosis*. ^11, 24^ Search for genes related to antibiotic and secondary metabolite pathways was performed by a deep analysis of protein sequences against the NCBI nr and KEGG databases. ^59^ Furthermore, we studied the natural product domain using CD-Search and NaPDos softwares. ^60, 61^ Phylogenetic trees were obtained using MEGA version 7 and the maximum likelihood (ML) algorithm, with 1,000 bootstrap replicates. ^53^ Phylogenetic reconstruction was analyzed using iTOL v3 online. ^62^ At last, we identified *A. castellanii* genes lmost closely related to ARMs homologous genes, and for each gene, a BLASTp search was performed against NCBI nr, with an e-value cutoff of 1e-03. ^51^ Deduced protein sequence alignments were carried out using Muscle software. ^52^ Phylogenetic trees were obtained using FastTree software and the maximum likelihood method with the Jones-Taylor-Thornton (JTT) model. Finally, we analyzed the phylogenetic trees using iTOL v3 online. ^62, 63^

### Genome annotation and analysis of *C. marseillensis*

Prodigal software was used for ORF prediction with the default settings. ^64^ Deviations in the sequencing regions predicted by ORFs were excluded. BLASTp was used to predict the bacterial proteins with an es-value cutoff of 1e-03, with a coverage of 70% and an identity of 30% according to the COG database. ^51, 55^ If there was no match, the BLASTp search in the database was extended with an e-value cutoff at 1e-03, coverage of 70% and an identity of 30%. ^46^ On the other hand, if the length of the sequence was less than 80 amino acids (aa), an e-value of 1e-05 was used. The HMMER web server was used in the domains that were maintained by the PFAM domains. ^65^ The rRNA and tRNA genes were retrieved using the RNAmmer and tRNAScanSE tools. ^66, 67^ ORFans were identified when the E value of BLASTp was less than 1e-03 for an alignment length greater than 80 amino acids. We also used the genome-to-genome distance calculator web service to calculate DNA: digital DNA hybridization estimates (dDDH) with confidence intervals according to recommended parameters (Formula 2, BLAST). ^68^

Genomic similarity of *C. marseillensis* with closely related species was estimated using OrthoANI software. ^69^ The pan-accessory genome distribution of *C. marseillensis* and other closely related species was evaluated using ClustAGE software. ^70^ The closely related species *Rickettsia bellii* (NC_007940), *Rickettsia conorii* (NC_003103.1), *Rickettsia heilongjiangensis* (NC_015866.1), *Rickettsia sibirica* (NZ_AABW01000001.1)*, Rickettsia parkeri* (NC_017044.1), *Rickettsia raoultii* (NZ_CP019435)*, Rickettsia rhipicephali* (NC_017042.1) and *Rickettsia typhi* (NC_006142) were used to perform genomic analysis using dDDH software, ClustAGE software and OrthoANI software.

## ACKNOWLEDGEMENTS

The authors acknowledge Morgan Gaia and Sandra Appelt for their contributive help in culturing the amoeba and paleomicrobiology studies, respectively. This work received (partial) support from Hitachi High-Tech Corporation.

## REFERENCES

1. Tito, R.Y., Knights, D., Metcalf, J., Obregon-Tito, A.J., Cleeland, L., Najar, F., Roe, B., Reinhard, K., Sobolik, K., Belknap, S., et al. (2012). Foster M, Spicer P, Knight R, Lewis CM Jr. Insights from characterizing extinct human gut microbiomes. PLoS One 7:e51146. doi: 10.1371/journal.pone.0051146. Epub 2012 Dec 12. PMID: 23251439; PMCID: PMC3521025.

2. 2. Reinhard, K.J., and Bryant, V.M. (1992). Coprolite analysis: a biological perspective on archaeology. Archaeological Method and Theory, Vol. 4, Michael B. Schiffer (ed.), pp. 245–288. © 1992 The University of Arizona Press, Tucson & London -ISSN 1043-1691.

3. 3. Appelt, S. 2013 Paléomicrobiologie des coprolithes. Thèse de doctorat, Aix-Marseille. See https://www.theses.fr/2013AIXM5068.

4. Malavin, S., and Shmakova, L. (2020). Isolates from ancient permafrost help to elucidate species boundaries in *Acanthamoeba castellanii* complex (Amoebozoa: Discosea). Eur. J. Protistol. 73, 125671. (doi:10.1016/j.ejop.2020.125671).

5. Sawyer, T.K., Visvesvara, G.S., and Harke, B.A. (1977). Pathogenic amoebas from brackish and ocean sediments, with a description of *Acanthamoeba hatchetti*, n. sp. Science 196, 1324–1325. (doi:10.1126/science.867031).

6. Davis, P.G., Caron, D.A., and Sieburth, JMcN. (1978). Oceanic amoebae from the North Atlantic: culture, distribution, and taxonomy. Trans. Am. Microsc. Soc. 97, 73–88. (doi:10.2307/3225685).

7. Schatz, S., Cometa, I., and Rogerson, A. (2011). Effect of microaerophilic environments on the growth of selected *Acanthamoeba* isolates. Invest. Ophthalmol. Vis. Sci. 52, 5806.

8. Haruta, S., and Kanno, N. (2015). Survivability of microbes in natural environments and their ecological impacts. Microbes Environ. 30, 123–125. (doi:10.1264/jsme2.ME3002rh).

9. Sriram, R., Shoff, M., Booton, G., Fuerst, P., and Visvesvara, G.S. (2008). Survival of Acanthamoeba cysts after desiccation for more than 20 years. J. Clin. Microbiol. 46, 4045– 4048. (doi:10.1128/JCM.01903-08).

10. Watson, S.P., Clements, M.O., and Foster, S.J. (1998). Characterization of the starvation-survival response of *Staphylococcus aureus*. J. Bacteriol. 180, 1750–1758.

11. Chen, T., He, L., Deng, W., and Xie, J. (2013). The *Mycobacterium* DosR regulon structure and diversity revealed by comparative genomic analysis. J. Cell. Biochem. 114, 1–6. (doi:10.1002/jcb.24302).

12. Schroeder, J.M., Booton, G.C., Hay, J., Niszl, I.A., Seal, D.V., Markus, M.B., Fuerst, P.A., and Byers, T.J. (2001). Use of subgenic 18S ribosomal DNA PCR and sequencing for genus and genotype identification of *Acanthamoebae* from humans with keratitis and from sewage sludge. J. Clin. Microbiol. 39, 1903–1911. (doi:10.1128/JCM.39.5.1903-1911.2001).

13. Clarke, M., Lohan, A.J., Liu, B., Lagkouvardos, I., Roy, S., Zafar, N., Bertelli, C., Schilde, C., Kianianmomeni, A., Bürglin, T.R., et al. (2013). Genome of Acanthamoeba castellanii highlights extensive lateral gene transfer and early evolution of tyrosine kinase signaling.

14. Genome Biol. *14*:R11. doi: 10.1186/gb-2013-14-2-r11. PMID: 23375108; PMCID: PMC4053784.

15. 14. Hasni, I., Andréani, J., Colson, P., and La Scola, B. (2020). Description of Virulent Factors and Horizontal Gene Transfers of Keratitis-Associated Amoeba Acanthamoeba Triangularis by Genome Analysis. Pathogens. 9:217. doi: 10.3390/pathogens9030217. PMID: 32188120; PMCID: PMC7157575.

15. Karlyshev, A.V. (2019). Remarkable features of mitochondrial DNA of *Acanthamoeba polyphaga* linc ap-1, revealed by whole-genome sequencing. Microbiol. Resour. Announc. 8, e00430–19. (doi:10.1128/MRA.00430-19).

16. Chelkha, N., Levasseur, A., Pontarotti, P., Raoult, D., Scola, B.L., and Colson, P. (2018). A phylogenomic study of *Acanthamoeba polyphaga* draft genome sequences suggests genetic exchanges with giant viruses. Front. Microbiol. 9, (doi:10.3389/fmicb.2018.02098).

17. Fritz-Laylin, L.K., Prochnik, S.E., Ginger, M.L., Dacks, J.B., Carpenter, M.L., Field, M.C., Kuo, A., Paredez, A., Chapman, J., Pham, J., et al. (2010). The genome of *Naegleria gruberi* illuminates early eukaryotic versatility. Cell 140, 631–642. doi: 10.1016/j.cell.2010.01.032. PMID: 20211133.

18. Hasni, I., Chelkha, N., Baptiste, E., Mameri, M.R., Lachuer, J., Plasson, F., Colson, P., and Scola, B.L. (2019). Investigation of potential pathogenicity of *Willaertia magna* by investigating the transfer of bacteria pathogenicity genes into its genome. Sci. Rep. 9, 1–12. (doi:10.1038/s41598-019-54580-6).

20. 19. Opperdoes, F.R., de Jonckheere, J.F., and Tielens, A.G.M. (2011). *Naegleria gruberi* metabolism. Int. J. Parasitol. 41, 915–924 (doi:10.1016/j.ijpara.2011.04.004).

20. Hug, L.A., Stechmann, A., and Roger, A.J. (2010). Phylogenetic distributions and histories of proteins involved in anaerobic pyruvate metabolism in Eukaryotes. Mol. Biol. Evol. 27, 311–324 (doi:10.1093/molbev/msp237).

21. Leger, M.M., Gawryluk, R.M.R., Gray, M.W., and Roger, A.J. (2013). Evidence for a hydrogenosomal-type anaerobic ATP generation pathway in *Acanthamoeba castellanii*. PloS One 8, e69532 (doi:10.1371/journal.pone.0069532).

22. Funo, K., Kitagawa, W., Tanaka, M., Sone, T., Asano, K., and Kamagata, Y. (2014). Draft genome sequence of *Tomitella biformata* AHU 1821T, isolated from a permafrost ice wedge in Alaska. Genome Announc. 2 (doi:10.1128/genomeA.00066-14).

23. Spyrou, M.A., Tukhbatova, R.I., Wang, C.C., Valtueña, A.A., Lankapalli, A.K., Kondrashin, V.V., Tsybin, V.A., Khokhlov, A., Kühnert, D., Herbig, A. et al. (2018). Analysis of 3800-year-old *Yersinia pestis* genomes suggests Bronze Age origin for bubonic plague. Nat Commun. 9:2234. doi: 10.1038/s41467-018-04550-9. PMID: 29884871; PMCID: PMC5993720.

24. Voskuil, M.I., Schnappinger, D., Visconti, K.C., Harrell, M.I., Dolganov, G.M., Sherman, D.R., and Schoolnik, G.K. (2003). Inhibition of respiration by nitric oxide induces a *Mycobacterium tuberculosis* dormancy program. J. Exp. Med. 198, 705–713. (doi:10.1084/jem.20030205).

25. Sun, C., Yang, G., Yuan, J., Peng, X., Zhang, C., Zhai, X., Luo, T., and Bao, L. (2017). *Mycobacterium tuberculosis* hypoxic response protein 1 (Hrp1) augments the pro-inflammatory response and enhances the survival of *Mycobacterium smegmatis* in murine macrophages. J. Med. Microbiol. 66, 1033–1044 (doi:10.1099/jmm.0.000511).

26. Yuan, Y., Crane, D.D., Simpson, R.M., Zhu, Y.Q., Hickey, M.J., Sherman, D.R., and Barry, C.E. (1998). The 16-kDa alpha-crystallin (Acr) protein of *Mycobacterium tuberculosis* is required for growth in macrophages. Proc. Natl. Acad. Sci. U. S. A. 95, 9578–9583 (doi:10.1073/pnas.95.16.9578).

27. Cole, S.T., and Barrell, B.G. (1998). Analysis of the genome of *Mycobacterium tuberculosis* H37Rv. Novartis Found. Symp. 217, 160–172; discussion 172–177 (doi: 10.1002/0470846526.ch12).

28. Zhang, Y., Deng, A., Zhang, H., Xi, F., Ying, L., Wang, J., and Bai, H. (2013). Epothilone O, a new member of this family from *Sorangium cellulosum* strain So0157-2. J. Antibiot. (Tokyo) 66, 285–286 (doi:10.1038/ja.2012.121).

29. Sander G. Mécanisme d’action du nouvel antibiotique kirromycine [Mechanism of action of the new antibiotic kirromycin]. Reprod Nutr Dev (1980). 21:185–192. French. PMID: 6130579.

30. Gershenzon, J., and Dudareva, N. (2007). The function of terpene natural products in the natural world. Nat. Chem. Biol. 3, 408–414 (doi:10.1038/nchembio.2007.5).

31. Mathew, B.B., Singh, H., Biju, V.G., and Krishnamurthy, N.B. (2017). Classification, source, and effect of environmental pollutants and their biodegradation. J. Environ. Pathol. Toxicol. Oncol. Off. Organ Int. Soc. Environ. Toxicol. Cancer 36, 55–71 (doi:10.1615/JEnvironPatholToxicolOncol.2017015804).

32. Greub, G., and Raoult, D. (2004). Microorganisms resistant to free-living amoebae. Clin. Microbiol. Rev. 17, 413–433 (doi:10.1128/CMR.17.2.413-433.2004).

33. Tokarz-Deptuła, B., Niedźwiedzka-Rystwej, P., Czupryńska, P., and Deptuła, W. (2019). Protozoal giant viruses: agents potentially infectious to humans and animals. Virus Genes 55, 574–591 (doi:10.1007/s11262-019-01684-w).

34. Sun, T-W., Yang, C-L., Kao, T-T., Wang, T-H., Lai, M-W., and Ku, C. (2020). Host range and coding potential of eukaryotic giant viruses. Viruses 12 (doi:10.3390/v12111337).

36. 35. Pagnier, I., Raoult, D., and La Scola, B. (2008). Isolation and identification of amoeba-resisting bacteria from water in human environment by using an *Acanthamoeba polyphaga* co-culture procedure. Environ. Microbiol. 10, 1135–1144 (doi:10.1111/j.1462-2920.2007.01530.x).

36. Krisko, A., and Radman, M. (2019). Protein damage, ageing and age-related diseases. Open Biol. 9, (doi:10.1098/rsob.180249).

37. Girod, M., Enjalbert, Q., Brunet, C., Antoine, R., Lemoine, J., Lukac, I., Radman, M., Krisko, A., and Dugourd, P. (2014). Structural basis of protein oxidation resistance: a lysozyme study. PloS One 9, e101642 (doi:10.1371/journal.pone.0101642).

38. Drancourt M, Raoult D. 2005 Palaeomicrobiology: current issues and perspectives. Nat. Rev. Microbiol. 3, 23–35. (doi:10.1038/nrmicro1063).

39. Hebsgaard MB, Phillips MJ, Willerslev E. 2005 Geologically ancient DNA: fact or artefact? Trends Microbiol. 13, 212–220. (doi:10.1016/j.tim.2005.03.010).

40. Appelt S, Armougom F, Le Bailly M, Robert C, Drancourt M. 2014 Polyphasic analysis of a Middle Ages coprolite microbiota, Belgium. PLoS ONE 9. (doi:10.1371/journal.pone.0088376).

41. Weekers PH, Wijen JP, Lomans BP, Vogels GD. 1996 Axenic mass cultivation of the free-living soil amoeba, Acanthamoeba castellanii in a laboratory fermentor. Antonie Van Leeuwenhoek 69, 317–322. (doi:10.1007/bf00399620).

42. Bou Khalil JY, Andreani J, Raoult D, La Scola B. 2016 A Rapid strategy for the isolation of new faustoviruses from environmental samples using *Vermamoeba vermiformis*. J. Vis. Exp. JoVE (doi:10.3791/54104).

43. Tito RY et al. 2008 Phylotyping and functional analysis of two ancient human microbiomes. PLoS ONE 3. (doi:10.1371/journal.pone.0003703).

44. Iñiguez AM, Reinhard K, Carvalho Gonçalves ML, Ferreira LF, Araújo A, Paulo Vicente AC. 2006 SL1 RNA gene recovery from *Enterobius vermicularis* ancient DNA in pre-Columbian human coprolites. Int. J. Parasitol. 36, 1419–1425. (doi:10.1016/j.ijpara.2006.07.005).

45. Raoult D, Aboudharam G, Crubézy E, Larrouy G, Ludes B, Drancourt M. 2000 Molecular identification by “suicide PCR” of Yersinia pestis as the agent of Medieval Black Death. Proc. Natl. Acad. Sci. 97, 12800–12803. (doi:10.1073/pnas.220225197).

46. Benson DA, Cavanaugh M, Clark K, Karsch-Mizrachi I, Ostell J, Pruitt KD, Sayers EW. 2018 GenBank. Nucleic Acids Res. 46, D41–D47. (doi:10.1093/nar/gkx1094).

47. Kim M-J, Park K-J, Ko I-J, Kim YM, Oh J-I. 2010 Different roles of DosS and DosT in the hypoxic adaptation of Mycobacteria. J. Bacteriol. 192, 4868–4875. (doi:10.1128/JB.00550-10).

48. Khelaifia S, Lagier JC, Nkamga VD, Guilhot E, Drancourt M, Raoult D. Aerobic culture of methanogenic archaea without an external source of hydrogen. Eur J Clin Microbiol Infect Dis. 2016;35(6):985–991. doi:10.1007/s10096-016-2627-7

49. Bolger AM, Lohse M, Usadel B. 2014 Trimmomatic: a flexible trimmer for Illumina sequence data. Bioinforma. Oxf. Engl. 30, 2114–2120. (doi:10.1093/bioinformatics/btu170)

50. Bankevich A et al. 2012 SPAdes: A new genome assembly algorithm and its applications to single-cell sequencing. J. Comput. Biol. 19, 455–477. (doi:10.1089/cmb.2012.0021)

51. Altschul SF. 2014 BLAST Algorithm. In *eLS*, American Cancer Society. (doi:10.1002/9780470015902.a0005253.pub2)

52. Edgar RC. 2004 MUSCLE: multiple sequence alignment with high accuracy and high throughput. Nucleic Acids Res. 32, 1792–1797. (doi:10.1093/nar/gkh340).

53. Kumar S, Stecher G, Tamura K. 2016 MEGA7: Molecular Evolutionary Genetics Analysis Version 7.0 for Bigger Datasets. Mol. Biol. Evol. 33, 1870–1874. (doi:10.1093/molbev/msw054).

54. Stanke M, Keller O, Gunduz I, Hayes A, Waack S, Morgenstern B. 2006 AUGUSTUS: ab initio prediction of alternative transcripts. Nucleic Acids Res. 34, W435–W439. (doi:10.1093/nar/gkl200).

55. Tatusov RL, Galperin MY, Natale DA, Koonin EV. 2000 The COG database: a tool for genome-scale analysis of protein functions and evolution. Nucleic Acids Res. 28, 33–36. (doi: 10.1093/nar/28.1.33).

56. Huerta-Cepas J, Forslund K, Coelho LP, Szklarczyk D, Jensen LJ, von Mering C, Bork P. 2017 Fast genome-wide functional annotation through orthology assignment by eggNOG-Mapper. Mol. Biol. Evol. 34, 2115–2122. (doi:10.1093/molbev/msx148).

57. Kanehisa M, Sato Y, Morishima K. 2016 BlastKOALA and GhostKOALA: KEGG tools for functional characterization of genome and metagenome sequences. J. Mol. Biol. 428, 726– 731. (doi:10.1016/j.jmb.2015.11.006).

58. Besemer J, Lomsadze A, Borodovsky M. 2001 GeneMarkS: a self-training method for prediction of gene starts in microbial genomes. Implications for finding sequence motifs in regulatory regions. Nucleic Acids Res. 29, 2607–2618. (doi: 10.1093/nar/29.12.2607)

59. Kanehisa M, Goto S. 2000 KEGG: Kyoto Encyclopedia of Genes and Genomes. Nucleic Acids Res. 28, 27–30. (doi: 10.1093/nar/28.1.27).

60. Marchler-Bauer A, Bryant SH. 2004 CD-Search: protein domain annotations on the fly. Nucleic Acids Res. 32, W327–W331. (doi:10.1093/nar/gkh454).

61. Ziemert N, Podell S, Penn K, Badger JH, Allen E, Jensen PR. 2012 The natural product domain seeker NaPDoS: a phylogeny based bioinformatic tool to classify secondary metabolite gene diversity. PloS One 7, e34064. (doi:10.1371/journal.pone.0034064).

62. Letunic I, Bork P. 2016 Interactive tree of life (iTOL) v3: an online tool for the display and annotation of phylogenetic and other trees. Nucleic Acids Res. 44, W242–W245. (doi:10.1093/nar/gkw290)

63. Price MN, Dehal PS, Arkin AP. 2009 FastTree: computing large minimum evolution trees with profiles instead of a distance matrix. Mol. Biol. Evol. 26, 1641–1650. (doi:10.1093/molbev/msp077)

64. Hyatt D, Chen G-L, LoCascio PF, Land ML, Larimer FW, Hauser LJ. 2010 Prodigal: prokaryotic gene recognition and translation initiation site identification. BMC Bioinformatics 11, 119. (doi:10.1186/1471-2105-11-119)

65. Potter SC, Luciani A, Eddy SR, Park Y, Lopez R, Finn RD. 2018 HMMER web server: 2018 update. Nucleic Acids Res. 46, W200–W204. (doi:10.1093/nar/gky448)

66. Lagesen K, Hallin P, Rødland EA, Stærfeldt H-H, Rognes T, Ussery DW. 2007 RNAmmer: consistent and rapid annotation of ribosomal RNA genes. Nucleic Acids Res. 35, 3100–3108. (doi:10.1093/nar/gkm160)

67. Lowe TM, Chan PP. 2016 tRNAscan-SE On-line: integrating search and context for analysis of transfer RNA genes. Nucleic Acids Res. 44, W54–57. (doi:10.1093/nar/gkw413)

68. Auch AF, von Jan M, Klenk H-P, Göker M. 2010 Digital DNA-DNA hybridization for microbial species delineation by means of genome-to-genome sequence comparison. Stand. Genomic Sci. 2, 117–134. (doi:10.4056/sigs.531120)

69. Lee I, Ouk Kim Y, Park S-C, Chun J. 2016 OrthoANI: An improved algorithm and software for calculating average nucleotide identity. Int. J. Syst. Evol. Microbiol. 66, 1100–1103. (doi:10.1099/ijsem.0.000760)

70. Ozer EA. 2018 ClustAGE: a tool for clustering and distribution analysis of bacterial accessory genomic elements. BMC Bioinformatics 19. (doi:10.1186/s12859-018-2154-x).

