## Supplementary material for "Resuscitating Sleeping Beauties: reviving a six-hundred-year-old amoeba and endosymbiont": Suplemental Table 1-Supplemental fig S1-S6

**Hasni Issam ^a,b^, Mamadou Lamine Tall ^a,b^, Matthieu Le Bailly ^c^, Philippe Colson ^a,b^, Anthony Levasseur ^a,b^, Didier Raoult ^a,b^, Bernard La Scola ^a,b^ and Michel Drancourt ^a,b^***

a. Aix-Marseille Univ., Institut de Recherche pour le Développement IRD 198, Assistance Publique – Hôpitaux de Marseille (AP-HM), Microbes, Evolution, Phylogeny and Infection (MEΦI), UM63.

b. Institut Hospitalo-Universitaire (IHU) - Méditerranée Infection, France.

c. Franche-Comté University, CNRS UMR 6249 Chrono-Environment, 25 030 Besançon, France.

* Corresponding author: Michel Drancourt

IHU Méditerranée Infection, 19-21 Bd Jean Moulin 13005 Marseille,

**SUPPLEMENTARY CONTENT**

**SUPPLEMENTARY TABLES**

**Tables S1- S6.**

**SUPPLEMENTARY FIGURES**

**Figures S1-S7.**

**Supplemental titles and legends of tables**

**Table S1: Amplicons generated by *ad hoc* PCR amplifications.**

**Table S2: Genome annotation of *A. castellanii strain* Namur.**

**Table S3:** a) hydrogen and carbon dioxide concentration measurements; b, c, d, e and f) genes related to dormancy and hypoxic metabolism. In b: Summary of gene related to survival, in c: Genes belonging *A. castellanii* strain Namur sharing homology with genes related to dormancy in *Mycobacterium tuberculosis*, in d: Genes related to anaerobic respiration in *Acanthamoeba castellanii* strain Namur, in e: Genes related to the *A. castellanii* anaerobic metabolism compared with *Tomitella biformata*, *Yersina pestis* and *Coprolita marseillensis* protein sequences, in f: Genes related to the *M. tuberculosis*

dormancy metabolism compared with *A. castellanii*, *Tomitella biformata*, *Yersina*

*pestis* and *Coprolita marseillensis* protein sequences.

**Table S4: Genes related to metabolism.** Gene belonging to

*Acanthamoeba castellanii* strain Namur involved in metabolism of Chloroalkane and

Chloroalkene (A), secondary metabolite (B), terpenoid backbone biosynthesis (C),

NRPS-PKS (D) and beta lactamase (E).

**Table S5: Horizontal gene transfers.** In a) genes of *Acanthamoeba*

*castellanii* sharing best hits with amoeba-resisting bacteria; In b) gene of

*Acanthamoeba castellanii* sharing best hits with giant viruses; In c) *A. castellanii*

protein sequences shared with amoeba resisting microorganisms related to different

clusters of gene categories.

**Table S6: Genome annotation of endosymbiont.**

**Table S1: Amplicons generated by *ad hoc* PCR amplifications^#^.**

| **ID** | **Amplification length (bp)** | **Hit description** | **E-value** | **Hit Accession ID** | **Percentage Coverage** | **Percentage ID** | **Environment of Hit Isolation** |
| --- | --- | --- | --- | --- | --- | --- | --- |
| Culture | 416 | *Acanthamoeba castellanii clone CF1-119b* | 0 | KC164234.1 | 99% | 98% | compost of composting facilities |
|  |  | *Acanthamoeba castellanii ATCC 50374* | 0 | U07413.1 | 99% | 98% | ---- |
| PCR Coprolite* | 323 | *Acanthamoeba sp. Had_008* | 4e-164 | FJ042636.1 | 100% | 99% | cornea (host: human/Israel) |
|  |  | *Acanthamoeba polyphaga isolate A8/SB2* | 4e-162 | GU596994.1 | 99% | 99% | environmental biofilm |
| PCR cysts * | 405 | *Acanthamoeba polyphaga Page-23* | 0 | AF019061.1 | 100% | 100% | ---- |

^#^ The amplicon identifier, its length (bp) and the annotation according to the best BLAST hits (BLASTn *versus* the nucleotide NCBI database) are summarized. The E-value, the hit accession identifier, the percent identity, the environment of described isolation and the reference are also provided. * Suicide PCR amplifications were performed; the primer pairs were used only in working areas, and no positive controls were incorporated.

**Figure S1**: **Culture of the endosymbiont in different hosts.** In a) (red arrow), endosymbiont cultivated within strain Namur; in b) (red arrow), endosymbiont cultivated within strain Neff. Pictures were obtained with an emission scanning electron microscope SU5000 (Hitachi, Japan). Scale bars are presented on the pictures.

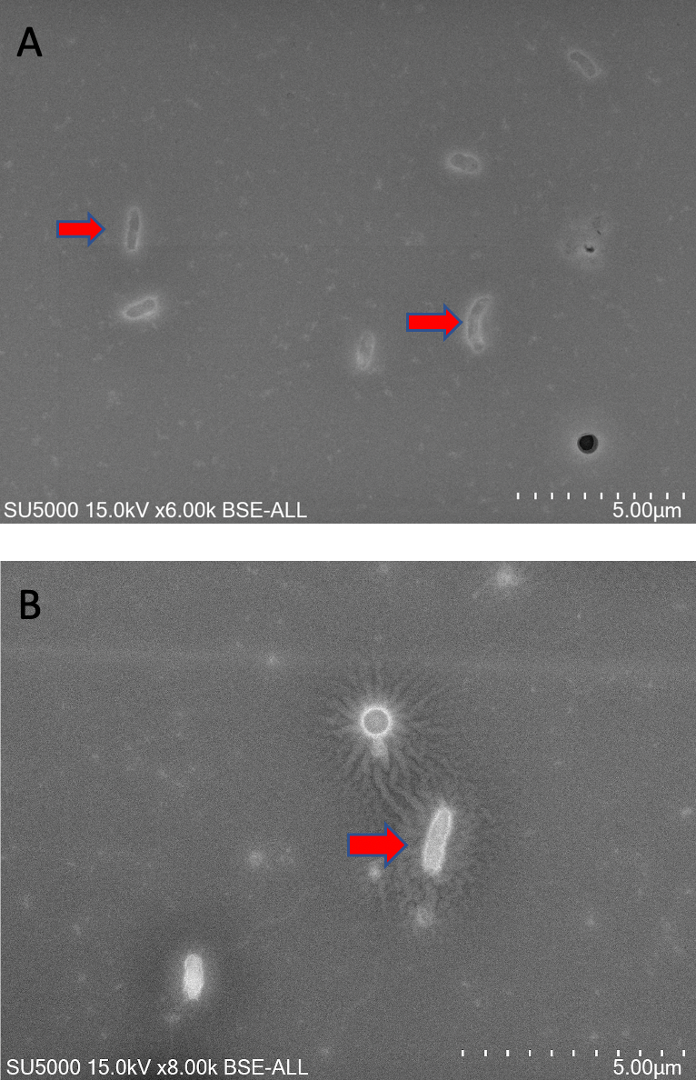

**Figure S2**: **Phylogenetic analysis of amoebas of the *A. castellanii* strain Namur.** The phylogenetic tree is based on partial available SSU rRNA sequences of amoebas. GenBank Accession numbers are indicated in parentheses. The sequences were aligned by Muscle, and trees were generated using the Jukes-Cantor model in MEGA 7.0.25 software. Numbers at the nodes are percentages of bootstrap values obtained by repeating the analysis 1,000 times to generate a consensus tree; only values with bootstraps ≥ 0.7 were displayed.

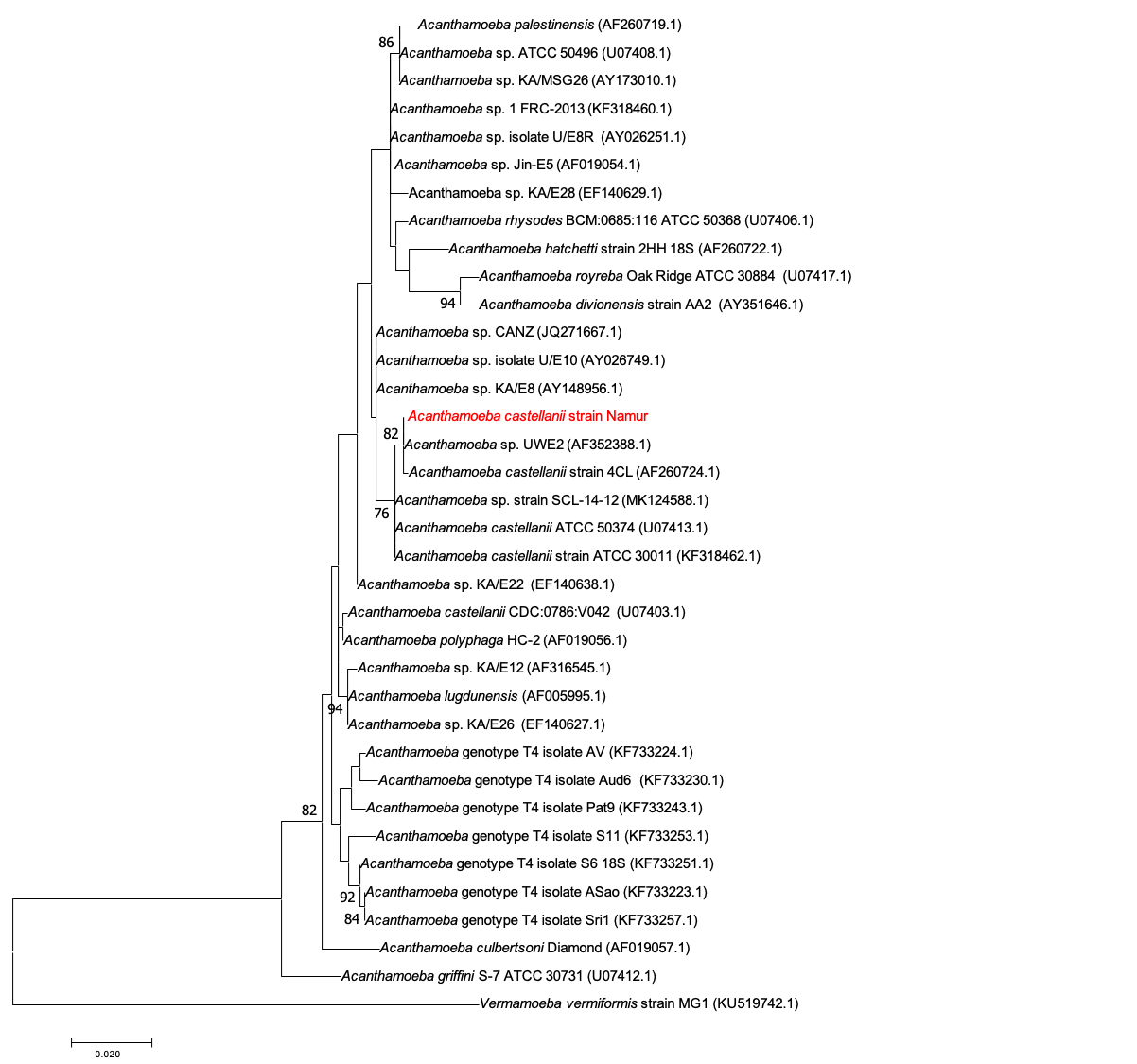

**Figure S3**: **Gene function of *A. castellanii* strain Namur.** In a) Representation of *A. castellanii* genes related to different clusters of gene categories; in b) distribution of *A. castellanii* protein sequences involved in different metabolic pathways. The *A. triangularis* protein was compared to the Kyoto Encyclopedia of Genes and Genomes Pathway (KEGG) database, and the representation of these protein sequences in the metabolic pathways was visualized.

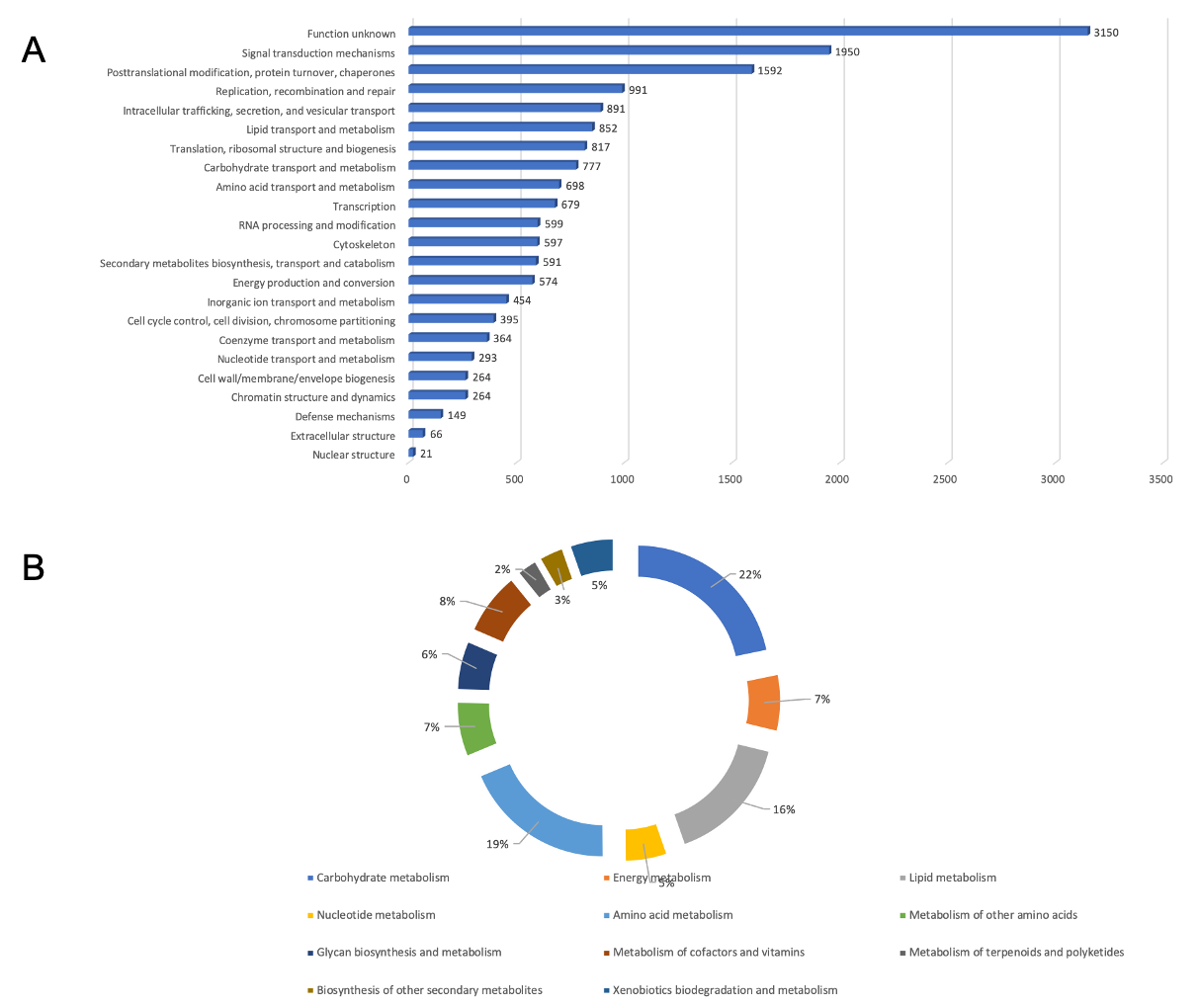

**Figure S4**: **Experimental progressive hypoxia.** Orange: Quantification of the production of H_2_, air (O_2_ and N) and CO_2_ at day 0. Black: Quantification of the production of H_2_, air and CO_2_ at day 14.

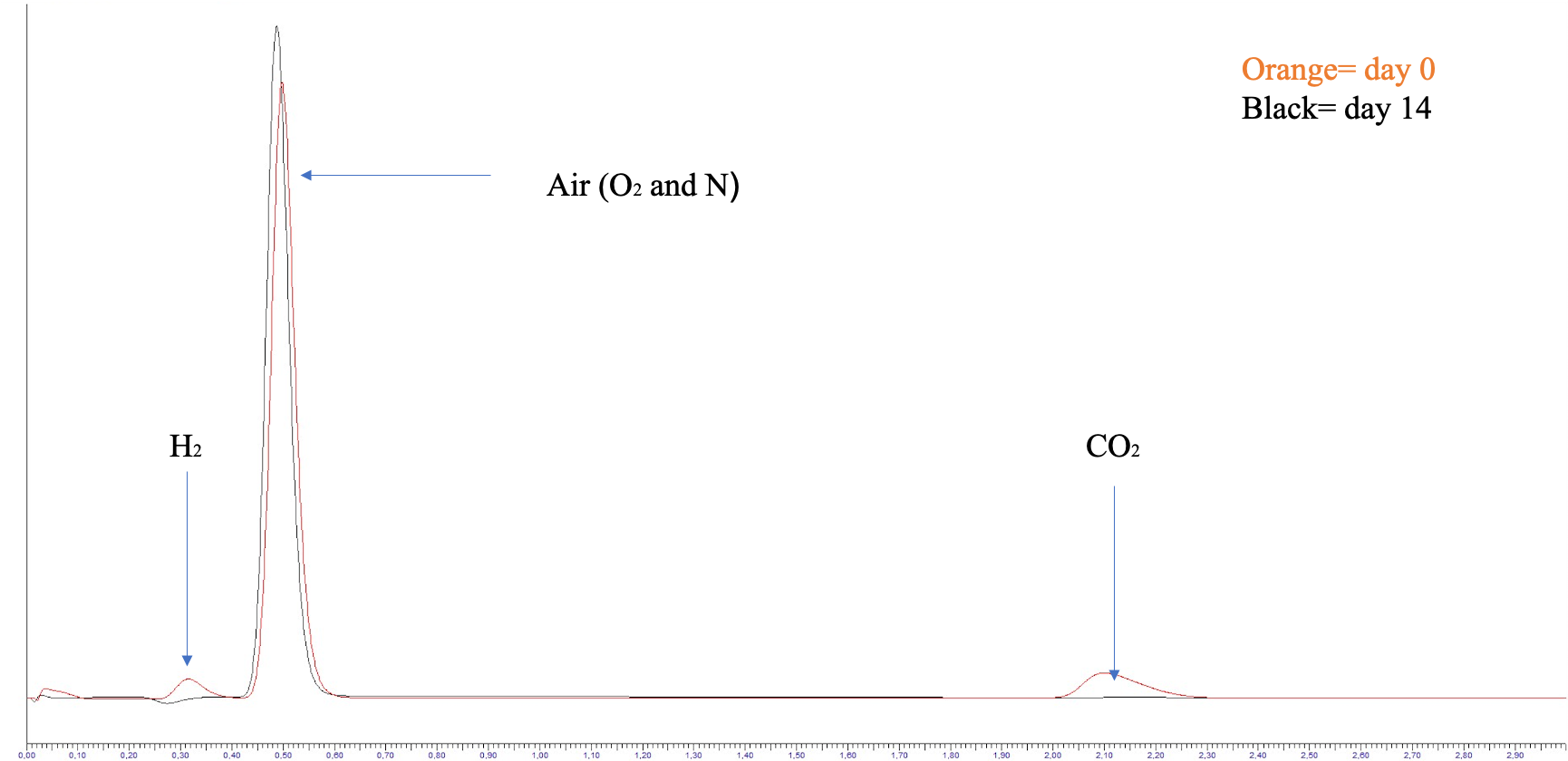

**Figure S5**: **Secondary metabolite analyze of *A. castellanii* strain Namur.** In a) Representation of conserved domain of beta-ketoacyl synthase. The analysis was performed by protein comparison against CDD database using the NCBI conserved domain website (<https://www.ncbi.nlm.nih.gov/Structure/cdd/wrpsb.cgi>). The characteristics of the conserved domain was reported as following: PksD: Acyl transferase domain in polyketide synthase (PKS) enzymes, FALL; FAAL belongs to the class I adenylate forming enzyme family and is homologous to fatty acyl-coenzyme A (CoA) ligases (FACLs), PKS_ER: Enoylreductase in Polyketide synthases. PKS_KR: This enzymatic domain is part of bacterial polyketide synthases. NADB-RO: A large family of proteins that share a Rossmann-fold NAD(P)H/NAD(P)(+) binding (NADB) domain. PKS_PP; Phosphopantetheine (or pantetheine 4' phosphate) is the prosthetic group of acyl carrier proteins (ACP) in some multienzyme complexes where it serves as a 'swinging arm' for the attachment of activated fatty acid and amino-acid groups. PP-binding: A 4'-phosphopantetheine prosthetic group is attached through a serine; In b) representation of lateral gene transfer analysis. Phylogenetic tree for *A. castellanii* proteins sharing homology with fungal proteins. The tree was constructed using the maximum-likelihood method based on the polyketide synthase sequences of *A. castellanii* strain Namur. The tree was constructed using 31 homologous sequences of *A. castellanii* retrieved by BLASTp on NCBI. In red, polyketide synthase sequences of *A. castellanii* strain Namur; in blue, homologs from *A. castellanii* strain Neff; in green, homologs from fungal organisms; in black, homologs from other organisms.

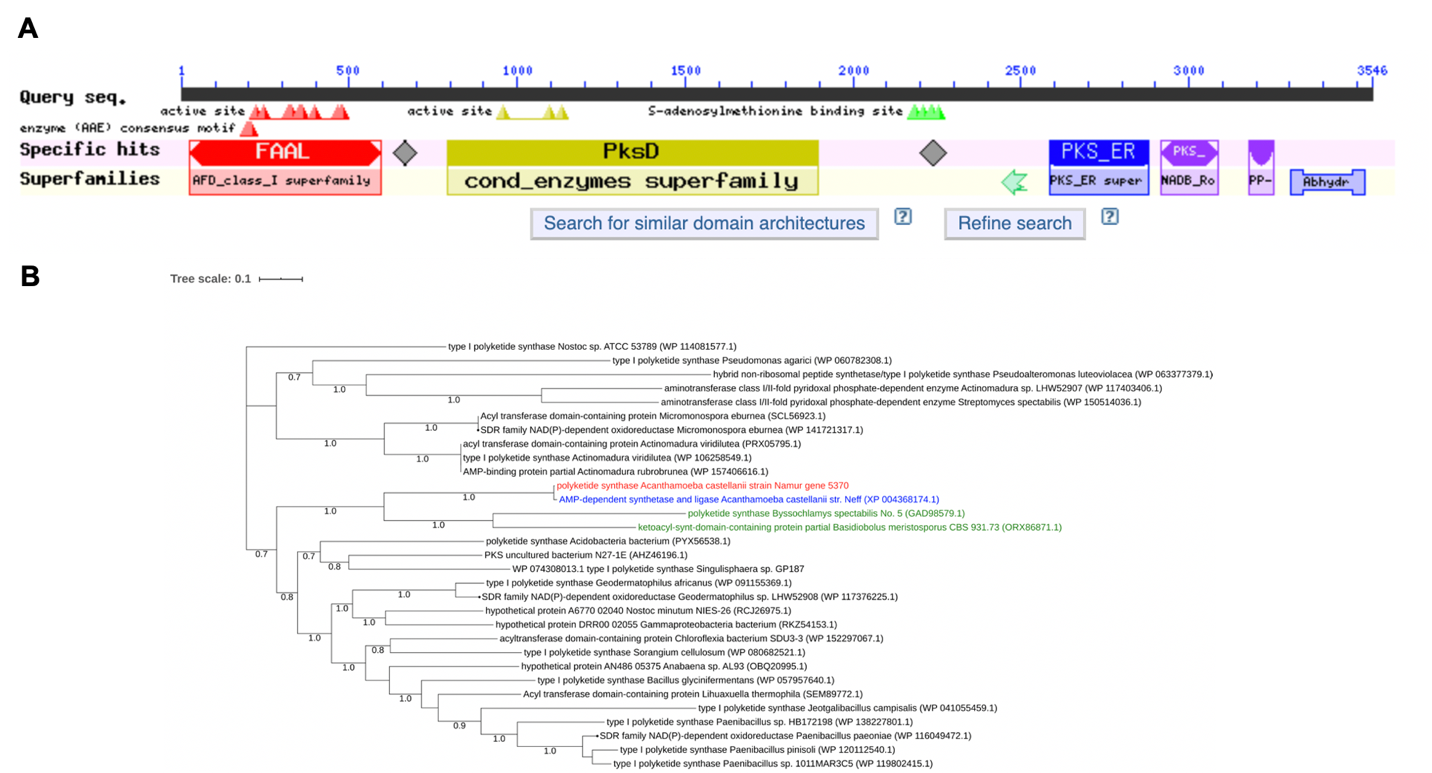

**Figure S6**: The comparison of COG annotation belonging to *C. marseillensis* and different closely phylogenetically related organisms.

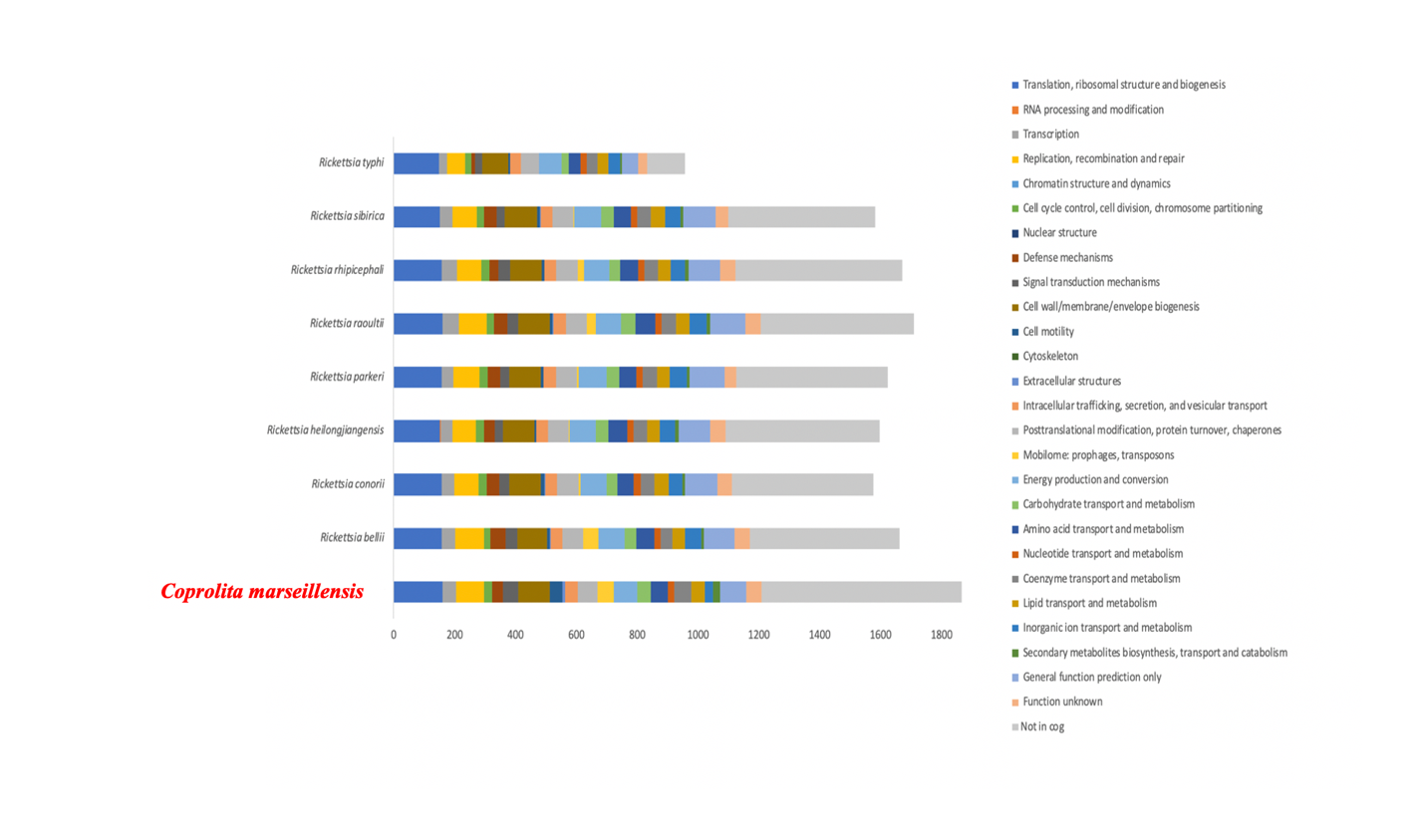

**Figure S7**: **Genomics analysis of endosymbiote.** In a) phylogenetic tree highlighting the position of *Coprolita marseillensis* with regard to other closely related species considering 16S rRNA. GenBank accession numbers of 16S rRNA are indicated in parentheses. Sequences were aligned using MUSCLE with default parameters, and phylogenetic inference was obtained using the maximum likelihood method and MEGA 7 software. Bootstrap values obtained by repeating the analysis 1,000 times to generate a majority consensus tree are indicated at the nodes. The scale bar indicates a 2% nucleotide sequence divergence; in b) Phylogenetic tree highlighting the position of *C. marseillensis* with regard to other closely related species based on whole genomes. The GenBank accession numbers of the genomes are indicated in parentheses. Sequences were aligned using MUSCLE with default parameters, and phylogenetic inference was obtained using the maximum likelihood method and MEGA 7 software. The bootstrap values obtained by repeating the analysis 1,000 times to generate a majority consensus tree are indicated at the nodes. The scale bar indicates a 10% nucleotide sequence divergence. In c) heatmap generated with OrthoANI values calculated using OAT software between *C. marseillensis* and other closely related species with standing in nomenclature.

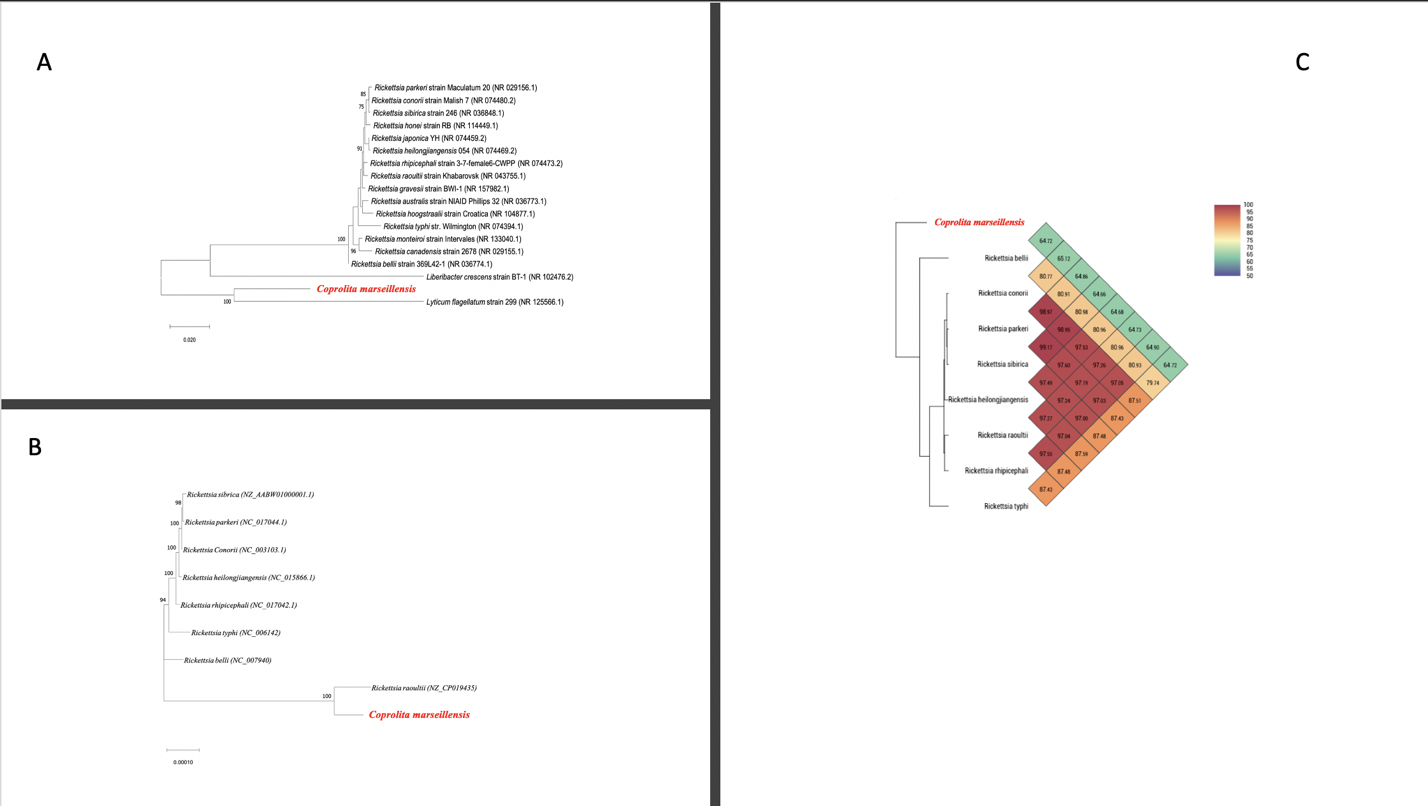
